# Epithelial Vegfa specifies a distinct endothelial population in the mouse lung

**DOI:** 10.1101/840033

**Authors:** Lisandra Vila Ellis, Margo P Cain, Vera Hutchison, Per Flodby, Edward D Crandall, Zea Borok, Bin Zhou, Edwin J Ostrin, Joshua D Wythe, Jichao Chen

**Author notes:** Contact: Jichao Chen.

## Abstract

The lung microvasculature is essential for gas exchange and commonly considered homogeneous. We show that *Vascular endothelial growth factor A* (*Vegfa*) from the epithelium specifies a distinct endothelial cell (EC) population in the postnatal mouse lung. *Vegfa* is predominantly expressed by alveolar type 1 (AT1) cells and locally required to specify a subset of ECs. Single cell RNA-seq identified 15-20% lung ECs as transcriptionally distinct and marked by Carbonic anhydrase 4 (Car4), which are specifically lost upon epithelial *Vegfa* deletion. Car4 ECs, unlike bulk ECs, have extensive cellular projections and are separated from AT1 cells by a limited basement membrane without intervening pericytes. Without Car4 ECs, the alveolar space is aberrantly enlarged despite the normal appearance of myofibroblasts. Lung Car4 ECs and retina tip ECs have common and distinct transcriptional profiles. These findings support a signaling role of AT1 cells and shed light on alveologenesis.

## INTRODUCTION

Endothelial cells (ECs) lining the blood vessels fulfill their transport function via region- and organ-specific specialization, such as the widely-known artery-capillary-vein relay and the non-leaky blood-brain barrier (Aird, 2007a, b; Potente and Makinen, 2017). Additional EC heterogeneity and plasticity are illustrated during development by the opposing duo of leading tip cells and trailing stalk cells within sprouting vessels, as well as the transition of ECs to hematopoietic, mesenchymal, and lymphatic lineages (Dejana et al., 2017; Gariano and Gardner, 2005). These functional and morphological differences in ECs are underlain by distinct gene expression profiles that have been extensively studied in the more tractable vasculature of the postnatal retina and, recently, have begun to be systematically tackled across organs using single cell RNA-seq (scRNA-seq) (Han et al., 2018; Sabbagh et al., 2018). An emerging theme for cell types that exist in multiple organs, as exemplified by macrophages (Lavin et al., 2014), is that they are endowed with organ-specific molecular signatures.

The pulmonary circulation consists of arterial and venous trees that parallel the branched airways and alveolar ducts and connect distally via a dense network of capillaries covering the gas-exchange alveoli – a high level of spatial coordination that presumably requires precise epithelial-endothelial crosstalk (Morrisey and Hogan, 2010). Although differences between lung macro- and micro-vasculature as well as lung-specific EC gene expression have been noted (Sabbagh et al., 2018; Stevens et al., 2008), the molecular, cellular, and genetic basis of these differences are poorly understood, especially in vivo (Durr et al., 2004). Deciphering lung EC heterogeneity and its developmental origin is also critical to our understanding of bronchopulmonary dysplasia, a severe lung disease often associated with premature birth and characterized by simplified alveoli and dysmorphic vasculature (Thebaud and Abman, 2007).

Our published work shows that (1) the lung capillaries are embedded within grooves of folded alveolar type 1 (AT1) cells, which constitute >95% of the alveolar epithelium; (2) developing AT1 cells, instead of alveolar type 2 (AT2) cells, express a potent angiogenic factor *Vascular endothelial growth factor A* (*Vegfa*); and (3) genetically blocking AT1 cell development decreases alveolar angiogenesis (Yang et al., 2016). These results indicate an instructive role of the alveolar epithelium for lung ECs. In this study, using scRNA-seq and single cell imaging, conditional and mosaic genetic models, and cross-organ comparisons, we found that epithelial-derived *Vegfa* is required to specify a transcriptionally-distinct lung EC population, which features net-like cellular extensions, embraces the epithelial contours, and promotes alveologenesis independent of myofibroblasts – contractile mesenchymal cells generally considered to drive alveologenesis (Bostrom et al., 1996).

## RESULTS

### Vegfa is predominantly expressed by AT1 cells and locally promotes alveolar angiogenesis

To confirm and extend our previous finding of *Vegfa* expression in developing AT1 cells (Yang et al., 2016), we immunostained developing and mature lungs carrying a nuclear LacZ knock-in reporter, *Vegfa^LacZ^* (Miquerol et al., 1999). While scattered and at a low level throughout the embryonic lung, the LacZ reporter in postnatal lungs co-localized with nuclei that were positive for NK2 homeobox 1 (NKX2.1; a lung epithelial lineage factor (Minoo et al., 1999)) but were not outlined by cuboidal E-Cadherin (a cell junction protein) staining – a characteristic feature of AT2 cells – indicating that AT1 cells, instead of AT2 cells, express *Vegfa* (Fig. 1A).

**Figure 1:**
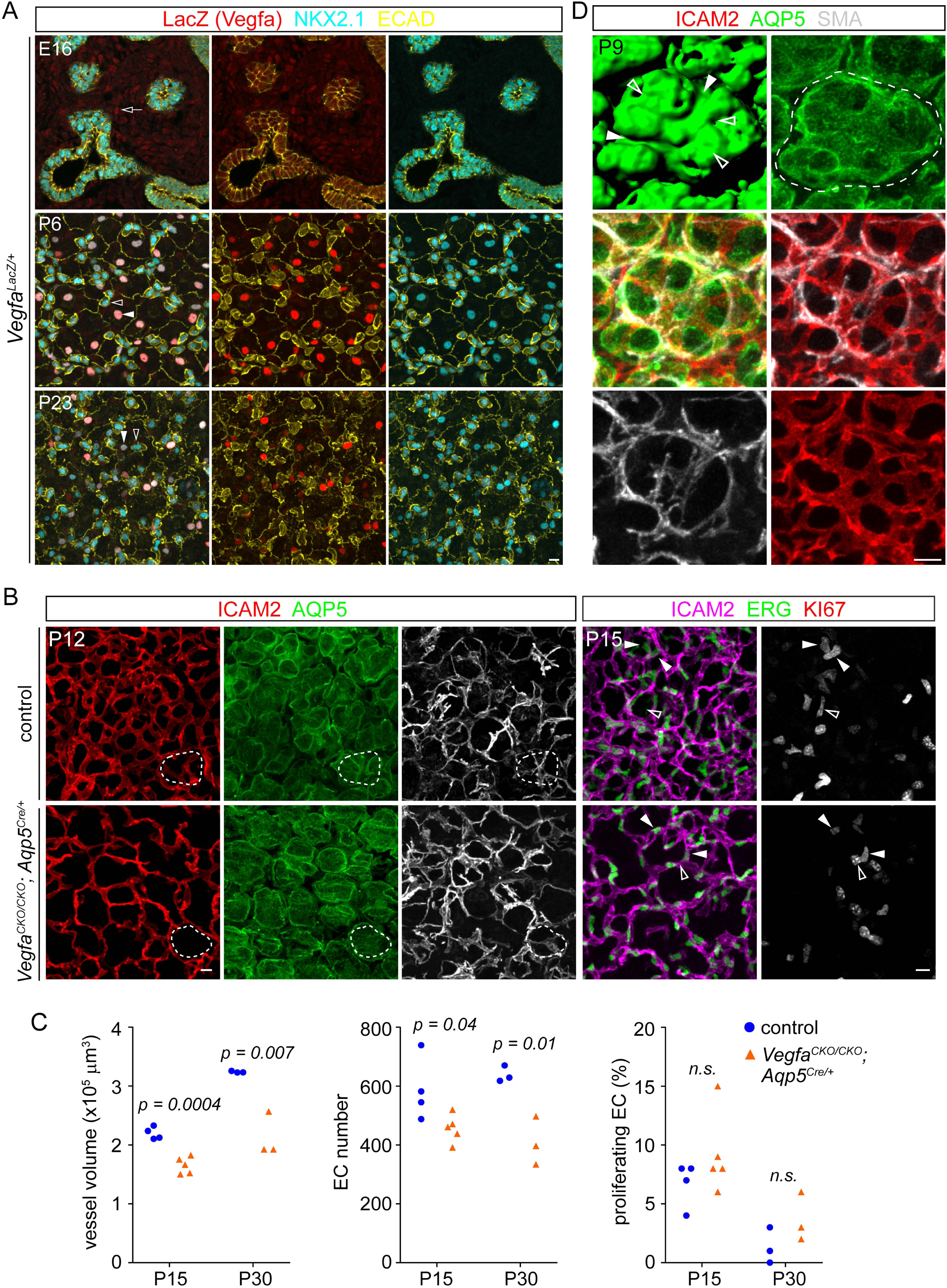
AT1 derived *Vegfa* is required for alveolar angiogenesis. See also Figure S1, S2. (A) Immunostained lungs with a nuclear LacZ knock-in allele of *Vegfa*, representative of at least 2 mice, showing its low mesenchymal expression at E16 (open arrow), and presence in AT1 cells (filled arrowhead) but absence in AT2 cells (open arrowhead). AT1 and AT2 cells are identified by NKX2-1 expression and distinct cell morphology, as outlined by E-Cadherin (ECAD). Postnatal lungs are shown as an en face view of the alveolar surface, which is better illustrated with surface rendering in (D). Scale: 10 um. (B) En face view of immunostained littermate lungs, representative of at least 3 littermate pairs, showing impaired alveolar angiogenesis in the *Vegfa* mutant. In the control lung, vessels (apical membrane marker ICAM2; nuclear marker ERG) cover alveolar islands (dash; AQP5) together with SMA-expressing myofibroblasts, whereas the remaining vessels in the mutant do not, despite normal coverage by myofibroblasts. Filled arrowhead, KI67/ERG double positive ECs. Non-ECs are also proliferative (open arrowhead). Scale: 10 um. (C) Quantification showing a lower vessel volume and EC number, but comparable proliferation (KI67^+^) in the mutant (Student’s t-test). Each symbol represents one mouse and is the average of three regions (318 um x 318 um x 20 um) with hundreds of EC cells counted for each region. (D) En face view of an immunostained alveolar island (dash) showing the epithelial surface (AQP5 rendering) with grooves containing both myofibroblasts (SMA) and vessels (ICAM2) (filled arrowhead) and those with only vessels (open arrowhead). Scale: 10 um.

To test the functional relevance of this AT1 specific *Vegfa* expression, we conditionally deleted *Vegfa* in AT1 cells using *Aqp5^Cre^* (Flodby et al., 2010), as validated by in situ hybridization probing for the deleted exon (Fig. S1A). The resulting mutant lungs had sparser alveolar vasculature without apparent changes in alpha-smooth muscle actin (SMA)-expressing myofibroblasts (Fig. 1B). This vascular phenotype was quantified as a persistent decrease in vessel volume and EC number (Fig. 1B, C). However, EC proliferation was not affected (Fig. 1B, C), suggesting that VEGFA does not function appreciably as a mitogen in the postnatal lung. Interestingly, the remaining vessels in the mutant were not randomly distributed: they failed to occupy the surface of alveolar “islands” that, in a control lung, contained grooves that were associated with myofibroblasts and thus considered secondary septation (Fig. 1B, D), raising the possibility that VEGFA is required for a specific subset of alveolar vessels.

The role of AT1-derived *Vegfa* was confirmed using another AT1 cell driver *Hopx^CreER^* (Takeda et al., 2011; Yang et al., 2016). A single dose of the chemical inducer, tamoxifen, resulted in mosaic recombination in AT1 cells, as indicated by the juxtaposition of recombined GFP^+^ and unrecombined GFP^-^ AT1 cells using a Cre reporter *Rosa^mTmG^* (Muzumdar et al., 2007)(Fig. 2A). As expected for the control lung, the alveolar islands, as defined in Fig. S1B, were invariably covered with vessels with no difference between GFP and non-GFP islands (Fig. 2). In contrast, up to 50% of the alveolar islands in the *Vegfa* mutant had no vessels – a phenotype more frequently observed for the GFP islands, presumably due to their higher likelihood of recombining both *Vegfa* alleles (Fig. 2). Notably, islands with and without vessel coverage were frequently found to be juxtaposed (Fig. 2 and Fig. S1B), indicating that AT1 derived VEGFA functions locally to promote vessel formation.

**Figure 2:**
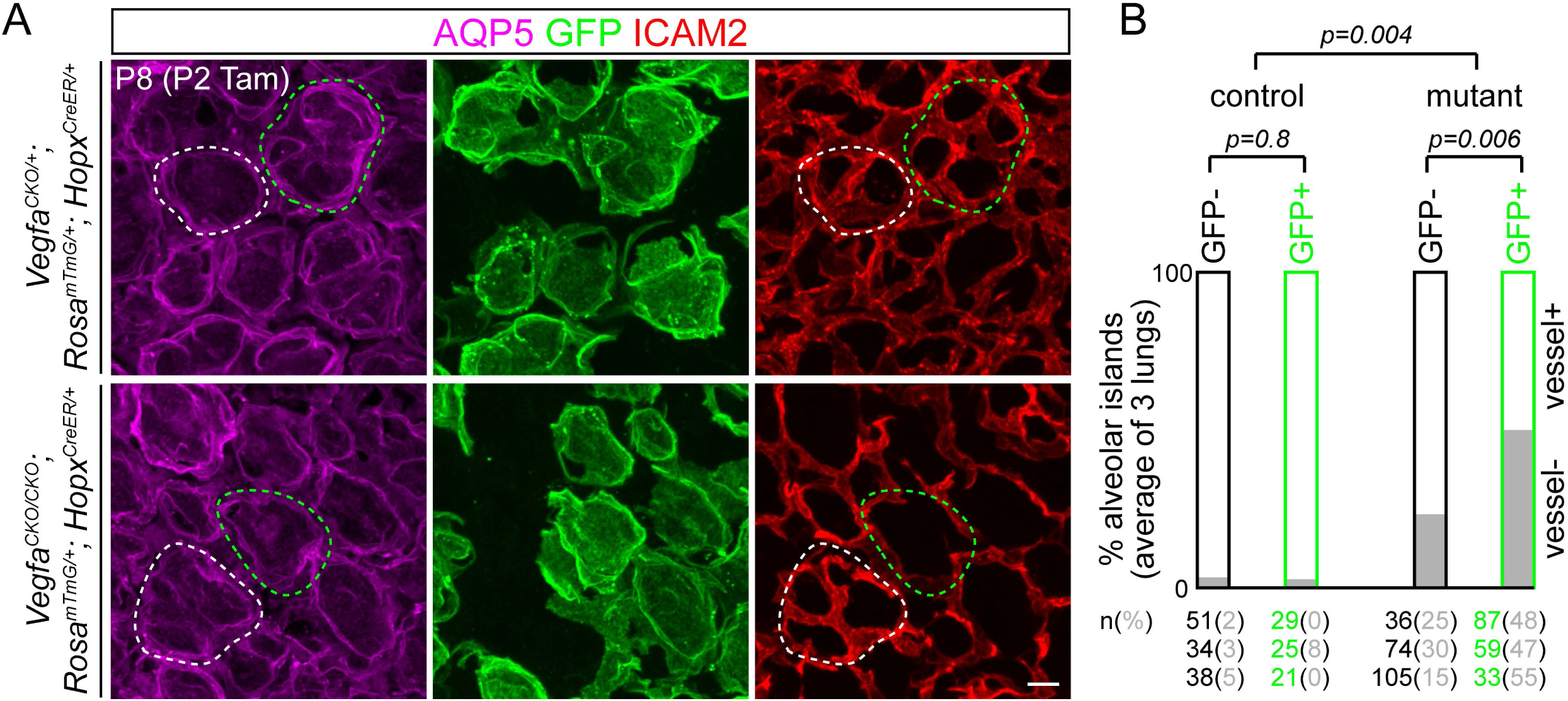
AT1 derived *Vegfa* promotes alveolar angiogenesis locally. See also Figure S1. (A) En face view of immunostained littermate lungs, representative of at least 3 littermate pairs, showing that, in the *Vegfa* mutant, juxtaposed recombined alveolar islands (GFP^+^; green dash) are associated with sparser vasculature than unrecombined ones (GFP^-^; white dash), whereas both types of alveolar islands have a comparable vasculature in the control, as expected. Tam, 200 ug tamoxifen. Scale: 10 um. (B) Quantification of vessel coverage in 3 littermate pairs. The number (n) of unrecombined (GFP^-^; black number) and recombined (GFP^+^; green number) alveolar islands examined in each mouse is tabulated with the percentage of aberrant islands (vessel^-^) in parenthesis (grey number), whose average is also shown as a stack bar graph. Although recombination of the *Rosa* locus does not always match that of the *Vegfa* locus, GFP^+^ islands are more likely to be aberrant (vessel^-^) in the mutant (Student’s t-test), which has more aberrant islands than the control (pooling unrecombined and recombined islands; Student’s t-test).

To exclude other functional sources of VEGFA, we conditionally deleted *Vegfa* from AT2 cells and ECs using *Sftpc^CreER^* and *Cdh5-CreER*, respectively (Barkauskas et al., 2013; Wang et al., 2010). As expected from the lack of appreciable expression (Fig. 1A), these mutants had normal vasculature (Fig. S1C, D).

Lastly, we noticed that the AT1-specific *Vegfa* mutant had normal vasculature at postnatal day (P) 2 (Fig. S2A), 5 days after the initiation of AT1 cell differentiation at embryonic day (E) 17 (Desai et al., 2014; Li et al., 2018a; Yang et al., 2016). This was likely because Cre recombination occurred after commitment to the AT1 identity – a requirement for a cell-specific Cre driver – such that enough VEGFA protein had accumulated and perdured after DNA deletion. To test this and to achieve more efficient deletion, we resorted to *Shh^Cre^*, which targets the entire epithelium during lung specification (Harris et al., 2006). Despite such early targeting, and different from reported results in another epithelial *Vegfa* deletion model (Yamamoto et al., 2007), branching morphogenesis was unaffected and vascular defects were only observed after E17, concomitant with AT1 cell differentiation (Fig. S2B, C), suggesting that early embryonic lung angiogenesis is supported by non-epithelial sources of *Vegfa* or *Vegfa*-independent mechanisms. The pan-epithelium *Vegfa* mutant recapitulated the AT1 specific *Vegfa* mutant phenotypes, including a decrease in vessel volume and EC number without affecting proliferation, as well as the failure to cover alveolar islands where secondary septation normally occurred (Fig. 3A and 3B). Taking into account all the cell type-specific *Vegfa* mutant models, we concluded that AT1 derived *Vegfa* is required locally for alveolar angiogenesis and possibly regulates a specific EC population in the developing lung.

**Figure 3:**
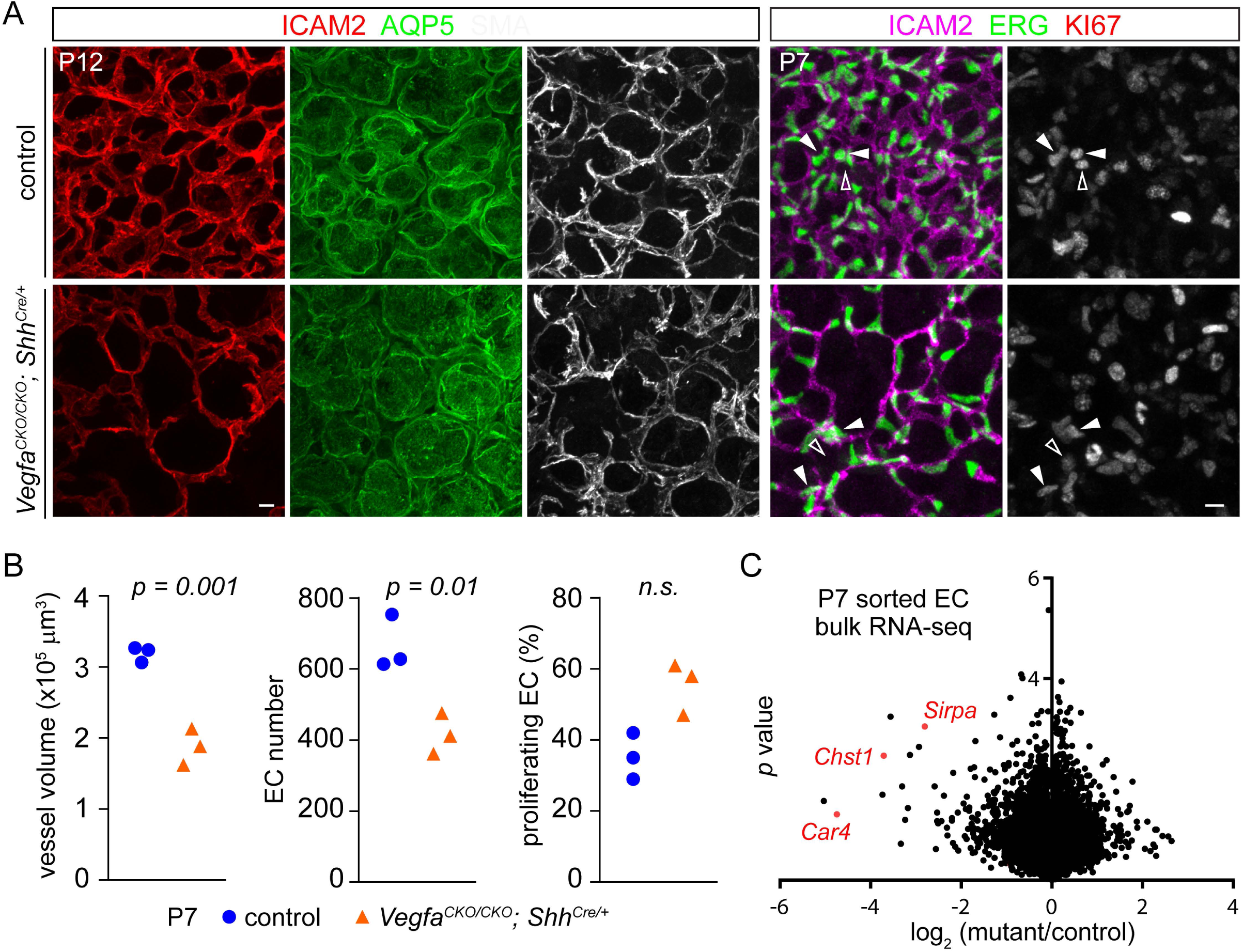
Epithelial *Vegfa* is required for alveolar angiogenesis. See also Figure S2, S3; Table S1. (A) En face view of immunostained littermate lungs, representative of at least 3 littermate pairs, showing impaired alveolar angiogenesis in the epithelial *Vegfa* mutant. Filled arrowhead, KI67/ERG double positive ECs. Non-ECs are also proliferative (open arrowhead). Scale: 10 um. (B) Quantification showing a lower vessel volume and EC number, but comparable proliferation (KI67^+^) in the mutant (Student’s t-test). Each symbol represents one mouse and is the average of three regions (318 um x 318 um x 20 um) with hundreds of EC cells counted for each region. (C) Volcano plot of bulk RNA-seq results from sorted lung ECs from 3 littermate pairs. Several downregulated genes are highlighted in red.

### Single cell RNA-seq identifies a molecularly distinct lung EC population

To understand the molecular basis of the *Vegfa* mutant phenotype, we optimized a cell dissociation and sorting protocol using lungs with genetically-labeled fluorescent ECs and found that adding a CD45 (an immune cell marker) negative selection step allowed better separation of ICAM2 (or CD31; both EC markers) positive and negative cells and that ICAM2 selection was consistent with, but more robust than, the more commonly-used CD31 selection (Fig. S3). Bulk RNA-seq comparison of purified ECs from control and epithelial *Vegfa* mutant lungs revealed downregulation of genes that were, intriguingly, recently identified as markers of sprouting tip ECs, such as S*ignal-regulatory protein alpha* (*Sirpa*) and *Carbohydrate sulfotransferase 1* (*Chst1*) (Fig. 3C and Table S1) (Sabbagh et al., 2018). This, together with the established role of *Vegfa* in inducing retinal tip ECs (Gariano and Gardner, 2005) and the non-random distribution of remaining ECs in our *Vegfa* mutant lungs, led us to hypothesize that *Vegfa* specifies a subset of ECs in the lung that are analogous to tip ECs in the retina.

To examine such potential lung EC heterogeneity, we performed scRNA-seq on 4,857 sorted P14 lung ECs (out of 5,175 cells) using 10x Genomics. Intriguingly, 14% of the sequenced ECs comprised a transcriptionally distinct cluster, which we named Car4 EC because *Carbonic anhydrase 4* (*Car4*) was its most specific marker (Fig. 4). All remaining ECs expressed *Plasmalemma vesicle associated protein* (*Plvap*) and included two transcriptionally distinct clusters – non-capillary ECs that expressed *Von Willebrand factor* (*Vwf*; thus named Vwf EC; 8.5% of all ECs) and lymphatic ECs that expressed *Prospero homeobox 1* (*Prox1*; 1.5% of all ECs) (Fig. 4A, B), both of which were confirmed to be spatially distinct from capillary ECs (Fig. S4). The remainder of the *Plvap*-expressing ECs (76% of all ECs) did not display a strong gene signature (Fig. 4C) – possibly because they were not as specialized as Car4 ECs, Vwf ECs, and lymphatic ECs – but were named Plvap EC for both simplicity and the availability of a PLVAP antibody for tissue localization. We note that *Car4*, *Plvap*, *Vwf*, and *Prox1* were used as population-specific markers, but not investigated for their functions in this study.

**Figure 4:**
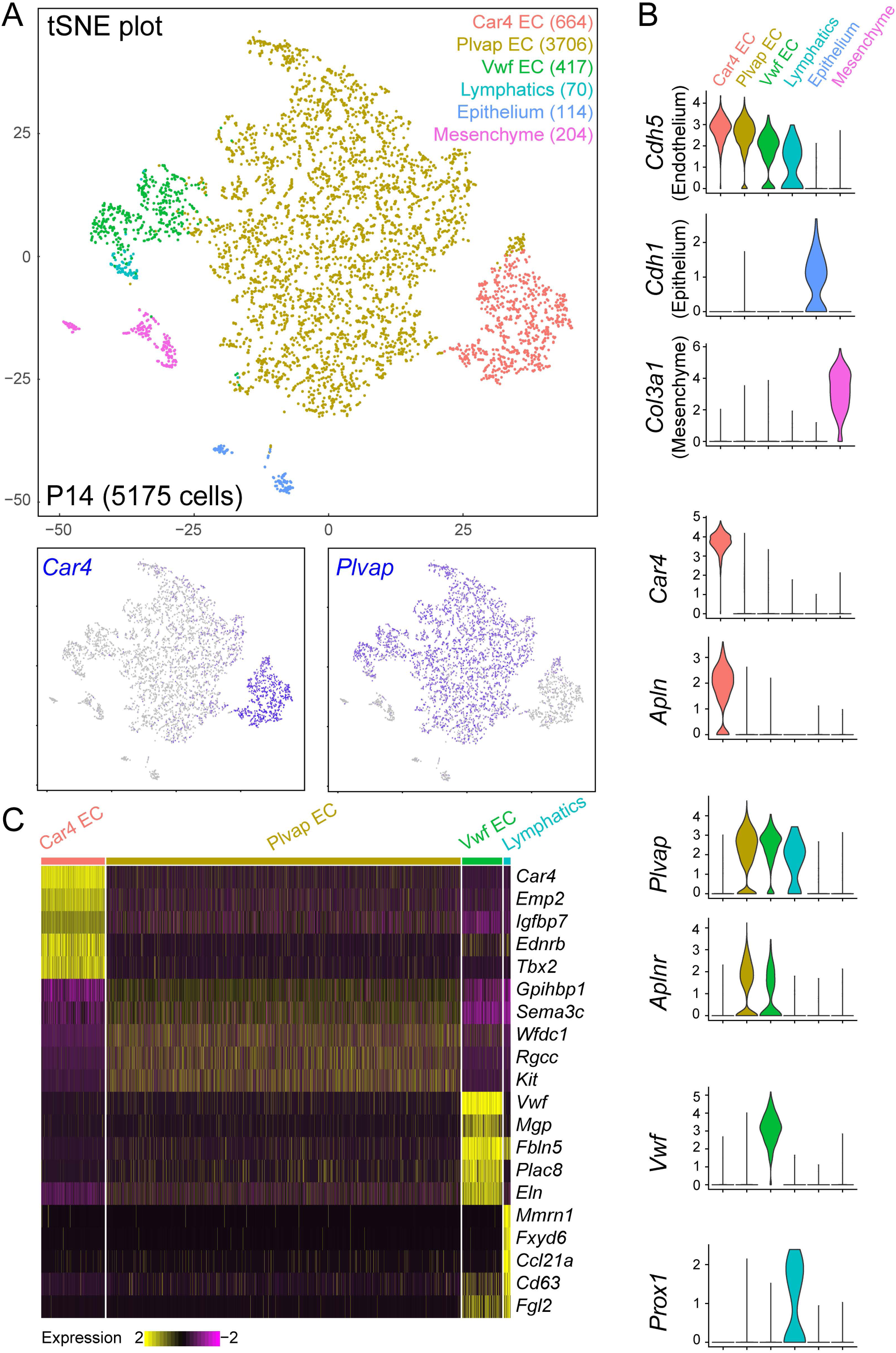
scRNA-seq identifies a distinct lung EC population. See also Figure S3, S4, S6, S7. (A) tSNE plot of scRNA-seq results from sorted P14 lung ECs with each cell population color-coded and the corresponding cell number shown in parenthesis. Lower panels: gene expression showing that *Car4* marks Car4 ECs; *Plvap* marks Plvap ECs, Vwf ECs, and lymphatic ECs. Epithelial and mesenchymal populations are minor contaminants from sorting. (B) Violin plots showing markers used to identify the six cell populations. (C) Heat map showing top 5 genes of each EC population. *Plvap* is expressed by all non-Car4 ECs, as shown in (A), and thus not among the top genes for Plvap ECs.

We further investigated the localization of Car4 and Plvap ECs by immunostaining in conjunction with a pan-EC nuclear marker ETS transcription factor (ERG) (Fish et al., 2017; Shah et al., 2016). Although predominantly on the cell membrane, both CAR4 and PLVAP also accumulated around the nucleus, allowing assignment of each ERG nucleus to the Car4 or Plvap EC population (Fig. 5A). Interestingly, CAR4 ECs covered the aforementioned alveolar islands that were undergoing secondary septation, whereas PLVAP ECs surrounded those islands, a distribution reminiscent of the remaining vessels in the *Vegfa* mutants (comparing Fig. 5A with Fig. 1B and 3A). This spatial difference in CAR4 and PLVAP staining was also evident for the corresponding nuclei (Fig. 5A). Furthermore, although CAR4 staining was abundant, the number of CAR4^+^ cells was under-represented (Fig. 4A, 5A), a discrepancy between cell number and vessel contribution suggesting that CAR4 ECs are disproportionally larger.

**Figure 5:**
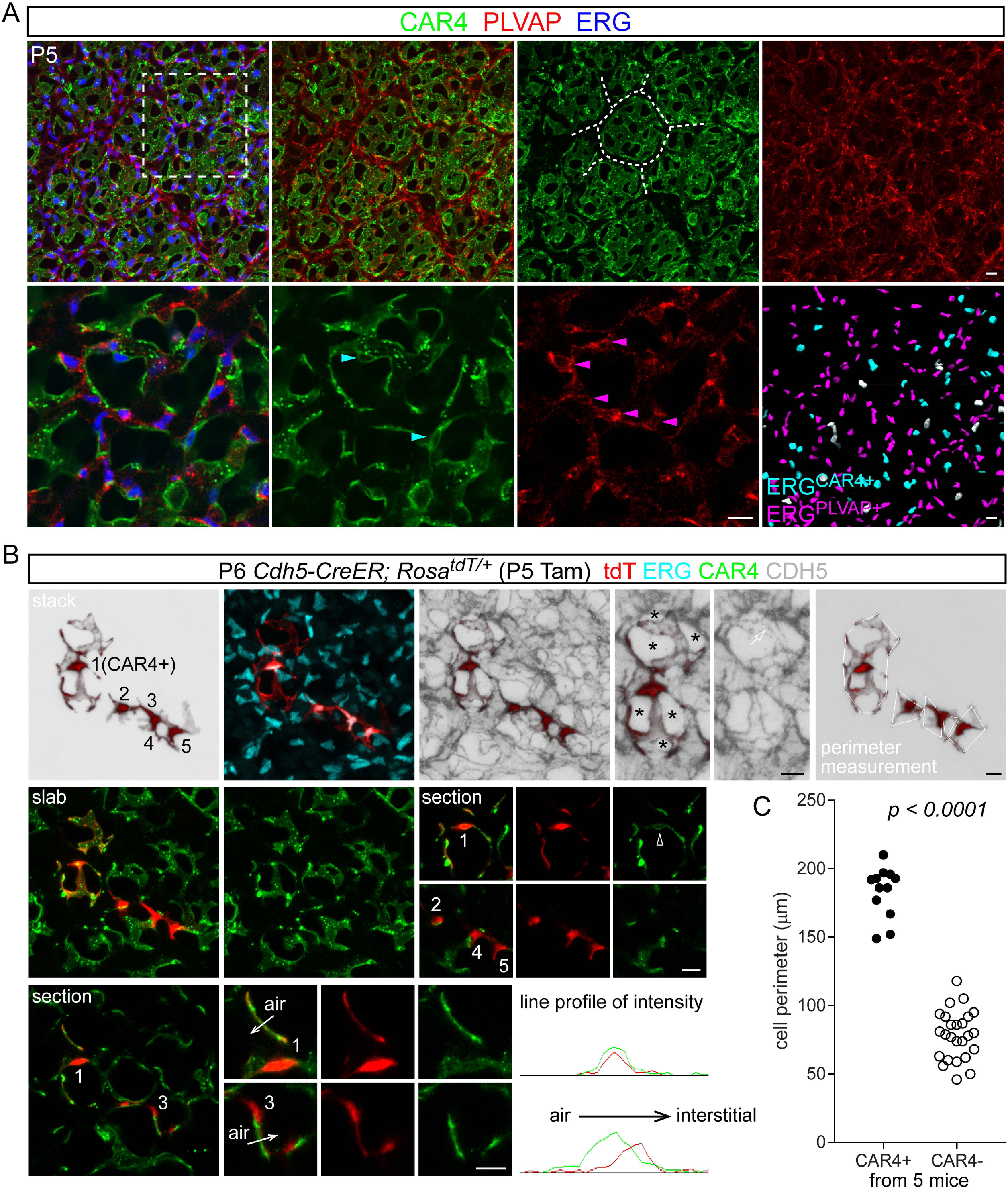
Car4 ECs have an expansive morphology. See also Figure S5, S6. (A) Representative en face view of immunostaining images from at least 5 mice. Boxed region is magnified as a section view in the first three images of the bottom row. CAR4 staining covers, whereas PLVAP staining surrounds, alveolar islands (dash). Perinuclear CAR4 and PLVAP staining allows assignment of ERG nuclei to CAR4 (cyan arrowhead) versus PLVAP (magenta arrowhead) ECs, which is automatically identified, as shown in the lower rightmost image (grey nuclei are ambiguous). Scale: 10 um. (B) Wholemount immunostaining of lungs with sparsely-labeled ECs, representative of at least 5 mice, viewed as a stack (40 um), a slab (top 20 um), or a section (1 um). Accumulation of tdT to ERG nuclei allows cell numeration (1 through 5). Cell #1 is a Car4 EC and cells #2-5 are non-Car4 ECs. Line profile analysis shows aligned versus shifted peaks for Car4 versus non-Car4 ECs, respectively. For shifted peaks, Car4 ECs are closer to the airspace than non-Car4 ECs (e.g. cell #3). Asterisk, avascular tissue surrounded by a single net-like Car4 EC. Open arrow, CDH5 junction overlapping with a single Car4 EC. Cell perimeter is measured by connecting protrusions that are visible in a projection view. Tam, 0.25 ug tamoxifen. Scale: 10 um. (C) Quantification of cell perimeter and comparison using Student’s t-test.

### Single cell imaging revealed extended Car4 EC morphology

Unlike the retina vasculature, which can be visualized in 2D after flat-mounting the tissue, lung ECs reside in thin tubes winding through a three-dimensional alveolar structure, making it challenging to examine their cell morphology. To overcome this, we adopted the sparse cell labeling method that we have used to study the similarly-complex AT1 cells (Yang et al., 2016). Specifically, we used a pan-endothelial inducible driver *Cdh5-CreER* (Wang et al., 2010), combined with an optimized, limited dose of tamoxifen, to sparsely label ECs so that individual ECs could be readily demarcated. As lungs were harvested within 24 hr after tamoxifen administration, labeled cells were not intended to be clonally related. We also used a strong reporter *Rosa^tdT^* (Madisen et al., 2010) to fill the entire cell body to (1) visualize the nucleus to confirm the intended single cell labeling and (2) colocalize with the membrane marker CAR4 to identify EC populations. Labeled cells were identified as Car4 ECs if they satisfied two criteria: (1) nuclear tdT completely overlapped with perinuclear CAR4 staining; (2) cytoplasmic tdT completely aligned with membrane CAR4 staining. Remarkably, in comparison to the bulk Plvap ECs, Car4 ECs were highly branched, often contributing to 5-10 vessel segments, and also much larger, as quantified by measuring the cell perimeter (Fig. 5B, C). Both Car4 and Plvap ECs had distinct morphology from the non-capillary Vwf ECs, which were much elongated along the direction of blood flow (Fig. S5A).

Closer examination of *Rosa^tdT^* labeled ECs in relation to neighboring Car4 ECs, an apical lumen marker Intercellular adhesion molecule 2 (ICAM2), and a cell adherens junction protein Cadherin 5 (CDH5) showed that (1) the sprawling labeled CAR4^+^ ECs did not exclude other ERG nuclei within those vessel segments; (2) the labeled CAR4^-^ ECs could be juxtaposed and share a common lumen with CAR4^+^ ECs, the latter being closer to the air space; (3) although the boundary of labeled cells coincided with CDH5, additional cell junctions were present within the labeled vessel segments (Fig. 5B and S5A). These observations suggested that individual lung capillaries are multi-cellular and can be comprised of both CAR4^+^ and CAR4^-^ ECs.

### Comparison of retina and lung ECs revealed common and organ-specific EC heterogeneity

The aforementioned similarity of Car4 ECs to tip ECs prompted us to examine tip EC-enriched genes in our distinct subsets of lung ECs. Intriguingly, Car4 ECs specifically expressed *Apln* as well as additional tip EC genes recently identified by scRNA-seq analysis of developing brain ECs (Sabbagh et al., 2018), such as *Plaur*, *Serpine1*, *Sirpa*, *Piezo2*, and *Chst1* (Fig. S6A, C). Plvap ECs specifically expressed several stalk cell genes including *Apelin receptor* (*Aplnr*) and *TEK receptor tyrosine kinase* (*Tek*; also known as *Tie2*) (Fig. S6A). However, the analogy of Car4/tip ECs and Plvap/stalk ECs did not extend to other known tip and stalk EC markers, as exemplified by *Esm1* and *Dll4* for tip ECs and *Hes1* and *Flt1* for stalk ECs (Blanco and Gerhardt, 2013) (Fig. S6A, B, C).

We further compared retina and lung ECs by immunostaining for ESM1, CAR4, and DLL4 (Fig. S6D). As reported (Rocha et al., 2014), ESM1 was restricted to tip ECs in the peripheral retina and excluded from mature vessels in the more central region. Interestingly, ESM1 was expressed in sporadic ECs near the lobe edge in embryonic lungs, raising the possibility that the lobe edge represents a growing front similar to the peripheral retina. However, this edge-specific expression was lost postnatally and ESM1 was enriched in ECs in a transition zone between capillaries and non-capillaries (Fig. S6D). In contrast, CAR4 was not detected in either the tip or stalk ECs in the retina, whereas its expression in the lung initiated at E19, concomitant with AT1 cell differentiation (Fig. S6D). Lastly, although *Dll4* regulates tip ECs (Blanco and Gerhardt, 2013), its protein was only slightly enriched in tip ECs and, in the central region, were more evident in arteries than veins. Similarly, DLL4 was widely expressed in embryonic lungs, but became more concentrated in a cord-like pattern postnatally (Fig. S6D). Such frequency and distribution of ESM1 and DLL4-expressing cells were also reflected in scRNA-seq (Fig. S6A, B). These data suggested that there are additional layers of EC heterogeneity in the lung that do not correspond to our four EC subpopulations.

Next, we examined the morphology of retina ECs in comparison to that of lung ECs using our sparse cell labeling method (Fig. S5B). In the central retina, ECs either were elongated along large vessels, similar to those in non-capillaries of the lung, or had a limited number of projections, similar to the lung Plvap ECs. In the peripheral retina, tip ECs displayed their characteristic blind-end filopodia, which was never found in the lung. Conversely, the expansive net-like morphology of Car4 ECs of the lung was also never found in the retina (Fig. S5B).

Among the tip EC genes shared by the lung Car4 ECs, we focused on *Apln* because of its reported role in sprouting angiogenesis (del Toro et al., 2010) and because its receptor *Aplnr* was specific to Plvap ECs (Fig. 4B). Using an *Apln^CreER^* knock-in, knock-out allele (Liu et al., 2015), we found that *Apln* underwent X-inactivation and, when tested with a *Rosa^L10GFP^* reporter, recombination was variable and enriched in but not specific to Car4 ECs (Fig. S7A, B). In addition, scRNA-seq comparison of sorted lung ECs from mutant (*Apln^CreER/Y^*) and littermate control males showed, as expected, a complete loss of *Apln* specifically in Car4 ECs, but no difference in either the number or gene expression of Car4 and Plvap ECs (Fig. S7C). Therefore, despite their intriguing expression patterns, the functional relevance of *Apln* and *Aplnr* in the lung is uncertain.

Taken together, the cross-organ comparison identified similarities and differences in EC gene expression and cell morphology, suggesting a common core vascular pathway superimposed with organ-specific adaptation.

### CAR4 ECs are specifically lost upon epithelial Vegfa deletion

Given their similarity to the retina tip ECs, which require *Vegfa* (Gerhardt et al., 2003), we asked if Car4 ECs of the lung also depended on *Vegfa*. To test this, we performed scRNA-seq on sorted ECs from control and epithelial *Vegfa* mutant lungs. Remarkably, the mutant lung specifically and completely lost the Car4 EC population, which represented 18% of all ECs in the control lung (Fig. 6A). In comparison, Plvap, Vwf, and lymphatic ECs were unaffected and proliferating ECs, marked by *Mki67* and frequent in early postnatal lungs, were also unaffected, consistent with our immunostaining-based quantification (Fig. 6A and 3B). This specific loss of Car4 ECs was confirmed by immunostaining showing that, in the mutant lung, CAR4 was rarely detected in the remaining vessels but unaffected in alveolar macrophages where CAR4 was normally present (Fig. 6A).

**Figure 6:**
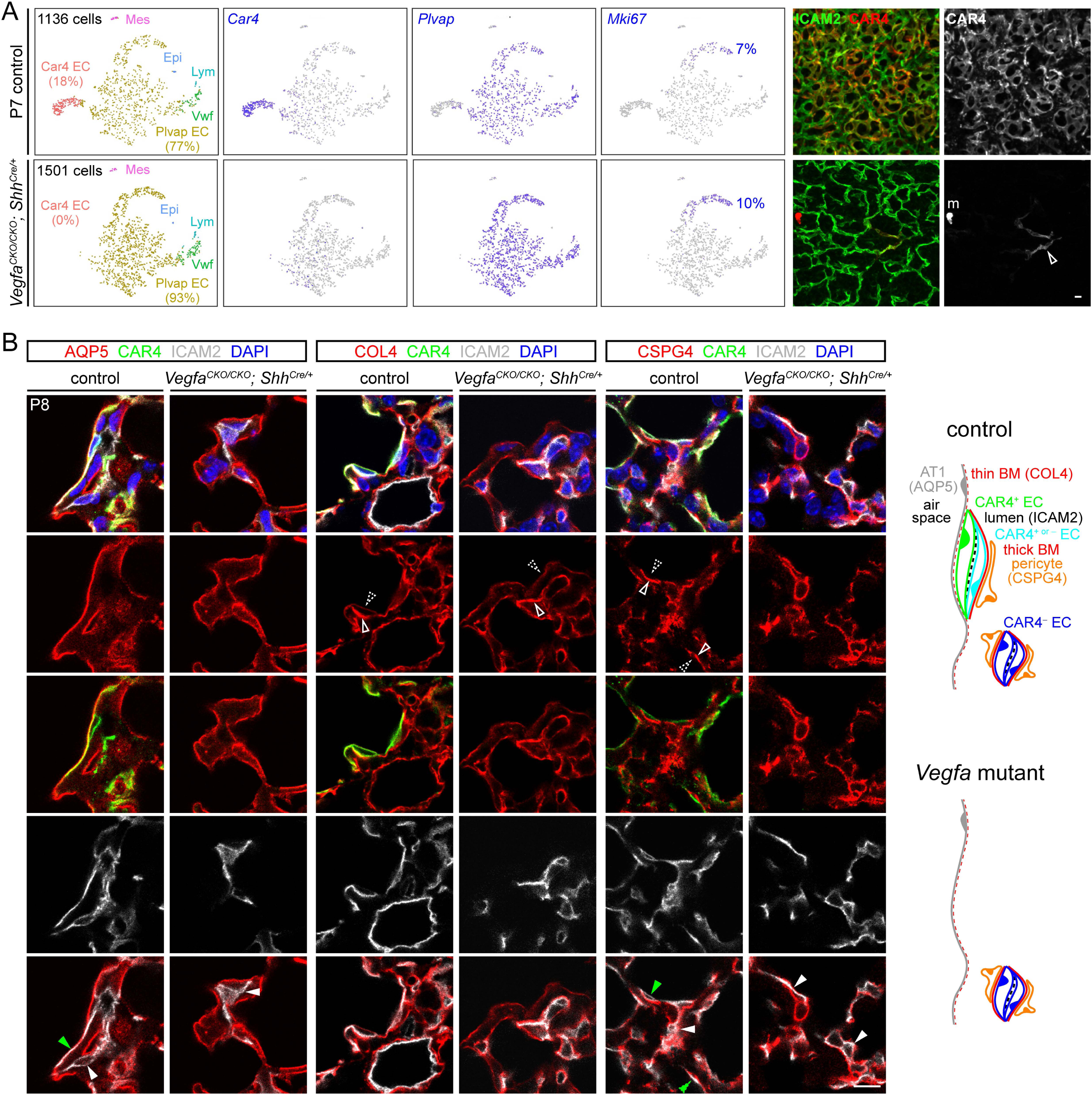
Car4 ECs are specifically lost upon epithelial *Vegfa* deletion. (A) Left panels: tSNE plot of scRNA-seq results from sorted ECs from littermate lungs showing a specific, complete loss of Car4 ECs in the mutant with the percentages out of all ECs in parenthesis. Plvap ECs and proliferating (*Mki67*) ECs are unaffected. Cell populations are color-coded as in Fig. 4A. Mes, mesenchyme; Epi, epithelium; Lym, lymphatic EC; Vwf, Vwf EC. Compared to P14 (Fig. 4A), P7 lungs have more proliferating (*Mki67*) ECs; their percentage is calculated with a UMI cutoff of 1. Right panels: en face view of immunostained littermate lungs, representative of at least 3 littermate pairs, showing rare Car4 staining in the remaining vessels in the mutant (open arrowhead). m, macrophage. Scale: 10 um. (B) Section immunostaining images of littermate lungs representative of at least 3 littermate pairs. As diagrammed, Car4 vessels (filled green arrowhead) abut the epithelium (AQP5), separated with a thin basement membrane (weak COL4 staining; dash open arrowhead) with no intervening pericytes (CSPG4; dash open arrowhead), but their sides away from the air space have a thicker basement membrane (strong COL4 staining; solid open arrowhead) and pericytes (solid open arrowhead). Non-Car4 vessels (filled white arrowhead; a subset of vessels in the control and all vessels in the mutant) do not abut the epithelium and are surrounded by a thick basement membrane and pericytes (solid open arrowhead). Scale: 10 um.

The specific loss of Car4 ECs in the *Vegfa* mutant lung allowed us to examine Car4 ECs at a higher resolution on sections and in the context of nearby non-ECs (Fig. 6B). Specifically, we used the pan-EC apical membrane marker ICAM2 to label the vessel lumen, which was mostly collapsed for capillaries, and used Aquaporin 5 (AQP5), Collagen type IV (COL4), and Chondroitin sulfate proteoglycan 4 (CSPG4; also known as NG2) to label the AT1 cell membrane, basement membrane, and pericyte membrane, respectively. We found that CAR4^+^ ECs – those that expressed CAR4 in the control but were absent in the *Vegfa* mutant – abutted the AT1 cell membrane and shared a thin basement membrane with the epithelium without intervening pericytes (Fig. 6B). In contrast, CAR4^-^ ECs – those that did not express CAR4 in the control or comprised all remaining vessels in the mutant – were away from AT1 cell membrane, underlain with a thicker basement membrane, and surrounded by pericyte processes (Fig. 6B). For vessels abutting the AT1 cell membrane, the side further away from the air space could consist of CAR4^-^ ECs, as described in Fig. 4B, and were found to be covered by a thicker basement membrane and pericyte processes (Fig. 6B and the diagram herein). The CAR4^-^ ECs in such hybrid vessels, as predicted by the lack of Car4 EC-containing vessels in the mutant, did not occupy the aforementioned secondary septae of the alveolar islands by themselves and depended on CAR4^+^ ECs for such locations (Fig. 3A, 6A), reminiscent of stalk ECs depending on tip ECs to spread to avascular regions. Taken together, these data suggested that Car4 ECs are induced by epithelial VEGFA, locate closest to the AT1 cell surface, and orchestrate the distribution of capillaries to secondary septae, as diagramed in Fig. 6B.

### CAR4 ECs contribute to alveolar morphogenesis independent of myofibroblasts

The CAR4^+^ vessels covering alveolar islands undergoing secondary septation and their specific absence in the *Vegfa* mutant (Fig. 1B, 3A and 6A) led us to examine the role of Car4 ECs in alveolar morphogenesis. Indeed, without the CAR4^+^ vessels coursing through the previously described grooves of the folding AT1 cells, the AT1 surface was smoother in the mutant (Fig. 7A). Consistent with a failure in secondary septation, the air space in the mutant lung was aberrantly enlarged with fewer recognizable alveoli (Fig. 7A), as quantified using a mean linear intercept method (Liu et al., 2017) as well as a D_2_ method that measured area-weighted air space diameters and bypassed the difficulty in distinguishing alveolar ducts and alveoli in the mouse lung (Jacob et al., 2009; Massaro and Massaro, 1996; Parameswaran et al., 2006). Notably, such defective alveologenesis occurred despite the normal specification and localization of myofibroblasts, a widely-accepted driver of secondary septation (Bostrom et al., 1996) (Fig. 7B).

**Figure 7:**
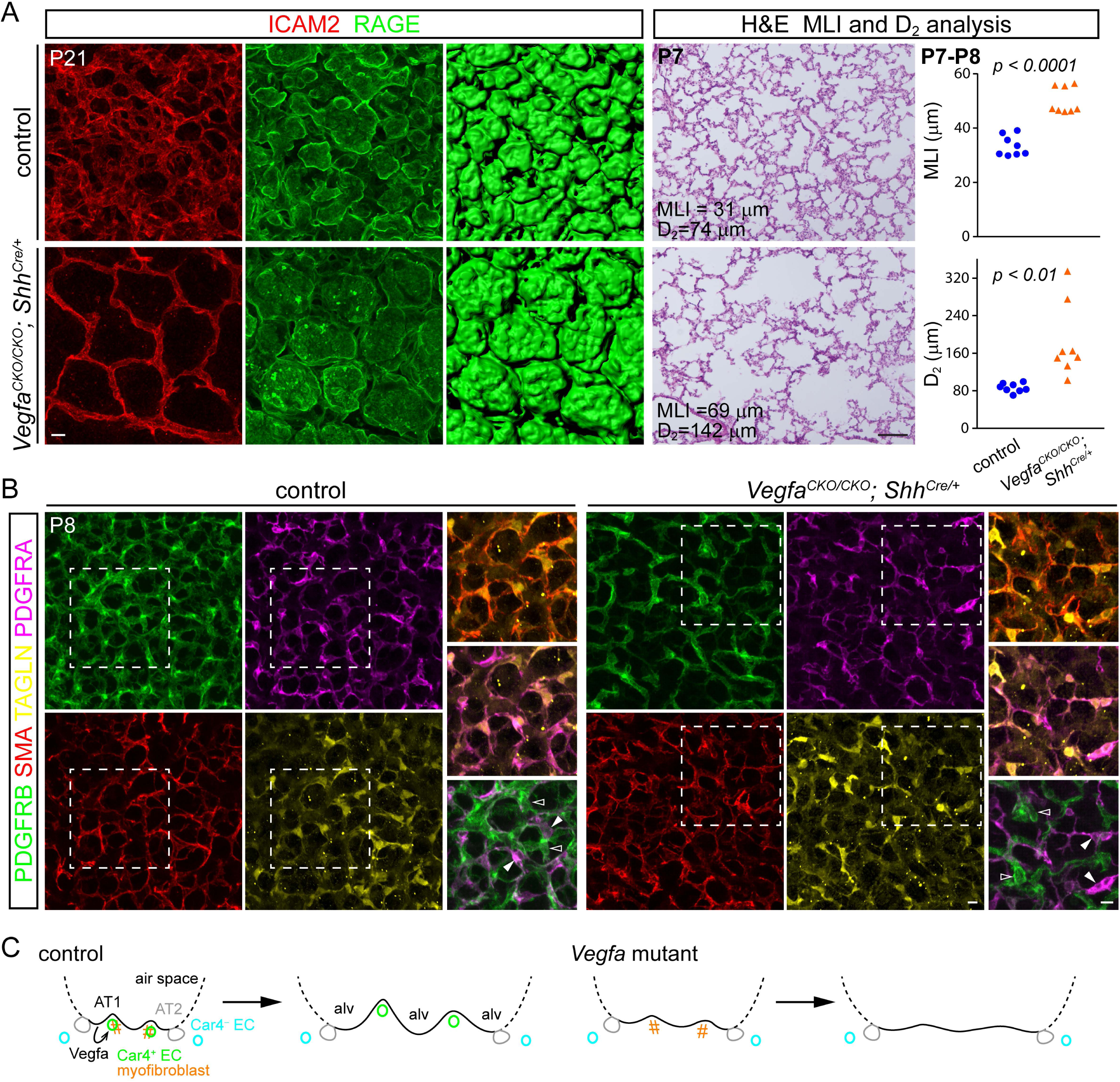
Aberrant alveolar enlargement in the absence of Car4 ECs but independent of myofibroblasts. (A) Left panels: en face view of immunostained littermate lungs, representative of at least 3 littermate pairs, showing a smoother surface (RAGE) of alveolar islands that are not subdivided by Car4 vessels in the mutant. Scale: 10 um. Right panels: H&E section images of littermate lungs with the corresponding mean linear intercept (MLI) and D_2_ measurements. Scale: 100 um. Each symbol represents one mouse and is the average of 3 regions (882 um x 664 um; Student’s t-test). (B) En face view of immunostained littermate lungs, representative of at least 3 littermate pairs, showing fewer pericytes (PDGFRB; open arrowhead) but normal myofibroblasts (SMA/TAGLN/PDGFRA triple positive, although variable in staining intensity; filled arrowhead) in the mutant. Boxed regions are magnified in merged views showing costaining of SMA, TAGLN, and PDGFRA that is distinct from PDGFRB. Scale: 10 um. (C) Schematics with color-coded cell types showing AT1 derived Vegfa signals to Car4 ECs, which, together with myofibroblasts, promote secondary septation and persist in the resulting septae even after disappearance of myofibroblasts in the mature lung. The *Vegfa* mutant fails to form Car4 ECs; non-Car4 ECs and myofibroblasts are insufficient for secondary septation, resulting in alveolar enlargement. Note that Car4 vessels may consist of Car4 ECs and non-Car4 ECs, as diagrammed in Fig. 6B. Alv, alveolus.

We further examined the mesenchymal lineage using a 4-marker panel: Platelet derived growth factor receptor, alpha polypeptide (PDGFRA), SMA, Transgelin (TAGLN; also known as SM22), and Platelet derived growth factor receptor, beta polypeptide (PDGFRB) (Fig. 7B). The peri-nuclear accumulation of PDGFRA, PDGFRB, and TAGLN allowed colocalization and assignment of cell types, while the commonly used myofibroblast marker SMA was not useful for such application but largely matched the cytoplasmic TAGLN staining. In the control lung, PDGFRB-expressing pericytes were distinct from PDGFRA cells, a subset of which expressed TAGLN and SMA and were thereby considered myofibroblasts. In the *Vegfa* mutant lung, there were fewer PDGFRB pericytes presumably as a result of loss of CAR4^+^ capillaries, but the PDGFRA/TALGN/SMA-expressing myofibroblasts were unaffected (Fig. 7B). These data are consistent with the notion that Car4 ECs, possibly together with the associated pericytes but independent of myofibroblasts, are required for alveolar morphogenesis.

## DISCUSSION

In this study, we show that the pulmonary microvasculature is heterogeneous and harbors a transcriptionally distinct EC population that is defined by CAR4 expression and specifically requires epithelial derived *Vegfa*, which is predominantly supplied by AT1 cells. These Car4 ECs feature an extended net-like morphology, situate seamlessly over the alveolar epithelium and specifically in regions undergoing secondary septation, and are required for alveolar morphogenesis independent of myofibroblasts (Fig. 7C). This work opens a new avenue of vascular research and has implications in alveolar development, physiology, and pathogenesis.

Our study reveals remarkable parallels between retinal tip ECs and lung Car4 ECs: (1) retinal astrocytes express *Vegfa* and provide the scaffold for tip ECs (Gariano and Gardner, 2005), whereas lung AT1 cells express *Vegfa* and provide the surface for Car4 ECs; (2) VEGFA in the retina induces tip ECs with characteristic filopodia and tip EC genes, whereas VEGFA in the lung induces Car4 ECs with characteristic net-like morphology and a subset of tip EC genes, such as *Apln*. However, there are also substantial differences: (1) the retina vasculature undergoes sprouting angiogenesis to spread from the central vascularized region to the peripheral avascular region in response to a hypoxia-induced VEGFA gradient, whereas the postnatal lung vasculature is surrounded by inhaled oxygen, always covers the alveolar epithelium as a dense net, and is believed to undergo intussusceptive angiogenesis (Burri et al., 2004) to match the expansion of the epithelial surface; (2) tip ECs disappear after development, but Car4 ECs persist even in the mature lung (Fig. S7C). Future mechanistic dissection of the common and distinct signaling events downstream of *Vegfa* in the two organs should shed light on the poorly understood process of intussusceptive angiogenesis and may possibly establish lung Car4 ECs as a novel model for vascular biology. More broadly, understanding organ-specific function of *Vegfa* may pave the way for targeted anti-VEGF therapy (Meadows and Hurwitz, 2012).

Alveologenesis, formation of alveoli, divides primary alveolar sacs – resulting from expansion of embryonic branch tips – into mature alveoli via the process of secondary septation (Yang and Chen, 2014). These secondary septae are marked by myofibroblasts, SMA-expressing contractile mesenchymal cells. Notably, we previously showed that such SMA-marked septae are the grooves of folded AT1 cells and coincide with capillaries that, as found in this study, are composed of Car4 ECs. Furthermore, although myofibroblasts are required for secondary septation in response to PDGFA signaling (Bostrom et al., 1996; Li et al., 2018b), our *Vegfa* mutant is missing Car4 ECs without affecting myofibroblasts or PDGFRA expression and yet displays aberrantly enlarged alveoli – consistent with a failure in secondary septation. This result raises the possibility that the force-generating myofibroblasts initiate secondary septation and AT1 cell folding, which is stabilized and maintained by CAR4^+^ vessels or their associated pericytes (Kato et al., 2018). This possibility is also consistent with the observation that myofibroblasts disappear or adopt other cell fates after the lung matures, whereas the vasculature persists (Li et al., 2018b; Yang et al., 2016). Future work will examine Car4 ECs in mutants that directly affect myofibroblasts and examine both myofibroblasts and Car4 ECs in experimental BPD models and patients with BPD, which is characterized by defective alveologenesis (Abman, 2001).

Car4 ECs are likely to have additional functions. First, compared to other ECs, Car4 ECs co-develop with the gas-exchanging AT1 cells, have a larger surface area, and are located closest to the alveolar epithelium, separated by a thinner basement membrane without intervening pericytes – all features suggesting a high efficiency in gas exchange. Testing this would require monitoring gas exchange in Car4^+^ versus Car4^-^ capillaries because conventional blood gas measurement of the *Vegfa* mutant will not distinguish a general loss of capillaries from a specific loss of high-efficiency Car4^+^ capillaries. Second, Car4 ECs specifically occupy the secondary septae and are required for alveolar morphogenesis, suggesting a structural role during lung development. Third, Car4 ECs specifically express secreted ligands including *Apln* and *Kitl* while the corresponding receptors, *Aplnr* and *Kit*, are expressed by Plvap ECs, suggesting possible signaling roles of Car4 ECs toward other ECs. However, our examination of the *Apln* mutant does not revealed any vascular phenotype. Future studies should focus on Car4 EC-specific genes, including CAR4 itself, which intriguingly is an enzyme involved in carbon dioxide formation (Crandall and O’Brasky, 1978; Fleming et al., 1993). Finally, the persistence of Car4 ECs in the mature lung calls for a better understanding of their role in homeostasis and injury-repair.

## ACKNOWLEDGEMENTS

We thank Drs. Napoleone Ferrara (Genentech; currently University of California San Diego, USA), Ralf Adams (University of Münster, Germany), Brigid Hogan (Duke University, USA) for providing the *Vegfa^CKO^*, *Cdh5-CreER*, and *Sftpc^CreER^* mice, respectively. We thank Kamryn Gerner-Mauro for assisting with scRNA-seq data analysis. The University of Texas MD Anderson Cancer Center DNA Analysis Facility and Flow Cytometry and Cellular Imaging Core Facility are supported by the Cancer Center Support Grant (CA #16672). This work was supported by the University of Texas MD Anderson Cancer Center Start-up Fund and National Institutes of Health R01-HL130129 (JC).

## AUTHOR CONTRIBUTIONS

LVE, JDW, and JC designed research; LVE, MPC, VH, EJO, and JC performed research; PF, EDC, and ZB provided the *Aqp5^Cre^* mice; BZ provided the *Apln^CreER^* mice; LVE, JDW, and JC wrote the paper; all authors read and approved the paper.

## METHODS

### Mice (*Mus musculus*)

The following mouse strains were used: *Vegfa^LacZ^* (Miquerol et al., 1999), *Vegfa^CKO^* (also called *VEGF-LoxP*) (Gerber et al., 1999), *Aqp5^Cre^* (Flodby et al., 2010), *Hopx^CreER^* (Takeda et al., 2011), *Sftpc^CreER^* (Barkauskas et al., 2013), *Cdh5-CreER* (Wang et al., 2010), *Apln^CreER^* (Liu et al., 2015), *Rosa^mTmG^* (Muzumdar et al., 2007), *Rosa^tdT^* (Madisen et al., 2010), *Rosa^L10GFP^* (Liu et al., 2014). The day of observing a vaginal plug was designated as E1. To induce Cre recombination, tamoxifen (T5648, Sigma) dissolved in corn oil (C8267, Sigma) was injected intraperitoneally. The tamoxifen dosage used is specified in the figure legends. Outliers were excluded only if there were technical errors, such as failed immunostaining. The number of control-mutant pairs and sections analyzed is stated in the figure legends. Unless specified, mice of both genders were used. Investigators were not blind to the genotypes. Control and mutant samples were processed in the same tube or block to minimize experimental variation. No power analysis was used to determine the sample size. All animal experiments were approved by the Institutional Animal Care and Use Committee at Texas A&M Health Science Center Institute of Biosciences and Technology and MD Anderson Cancer Center.

### Antibodies

The following antibodies were used: rabbit anti-Aquaporin 5 (AQP5, 1:2500, ab78486, Abcam), goat anti-Carbonic anhydrase IV (CAR4, 1:500, AF2414, R&D), BV786 rat anti-CD31 (1:250, 740870, BD Biosciences), PE/Cy7 rat anti-CD45 (1:250, 103114, BioLegend), mouse anti-Claudin 5 (Cldn5, 1:500, Invitrogen, 352588), rabbit anti-collagen IV (COL4, 1:2500, LSL-LB-1403, CosmoBioUSA), goat anti-Delta like canonical Notch ligand 4 (DLL4, 1:250, AF1389, R&D), Alexa Fluor 488 rat anti-CD324 (ECAD, 1:500, 53-3249-80, eBioscience), goat anti-Endothelial cell specific molecule 1 (ESM1, 1:500, AF1999, R&D), rabbit anti-Avian erythroblastosis virus E-26 (v-ets) oncogene related (ERG, 1:5000, ab92513, Abcam), goat anti-Vegfr3/Flt4 (1:1000, R&D, AF743), chicken anti-beta Galactosidase (LacZ, 1:500, Ab9361, Abcam), chicken anti-Green fluorescent protein (GFP, 1:5000, AB13970, Abcam), Alexa Fluor 647 rat anti-Intercellular adhesion molecule 2 (ICAM2, 1:500, A15452, ThermoFisher), rat anti-Intercellular adhesion molecule 2 (ICAM2, 1:2500, 16-1021-82, eBioscience), goat anti-Intercellular adhesion molecule 2 (ICAM2, 1:500, AF774, R&D systems), eFluor 570 rat anti-Ki67 (1:500, 41-5698-82, eBioscience), rabbit anti-Ki67 (1:1000, RM9106S0, ThermoFisher), rabbit anti-Chondroitin sulfate proteoglycan 4 (CSPG4, 1:1000, AB5320, Millipore), rabbit anti-NK2 Homeobox 1 (NKX2.1, 1:1000, sc-13040, Santa Cruz), rat anti-Platelet derived growth factor receptor alpha (PDGFRA, 1:1000, 14-1401-82, eBioscience), goat anti-Platelet derived growth factor receptor beta (PDGFRB, 1:1000, AF1042, R&D systems), rat anti-Plasmalemma vesicle associated protein (PLVAP, 1:125, 553849, BD Biosciences), rabbit anti-Prospero Homeobox 1 (PROX1, 1:250, 11-002, AngioBio), rat anti-Advanced glycosylation end-product specific receptor (RAGE, 1:1000, MAB1179, R&D systems), rabbit anti-Red fluorescent protein (RFP, 1:1000, 600-401-379, Rockland), Cy3-conjugated mouse anti-alpha-Smooth muscle actin (SMA, 1:1000, C6198, Sigma), rabbit anti-SM22 (TAGLN, 1:2500, Abcam, ab14106), Alexa Fluor 647 rat anti-Vascular endothelial cadherin (VECAD/CDH5, 1:250, 562242, BD Biosciences), rabbit anti-Von Willebrand Factor (VWF, 1:2500, Abcam, ab6994).

### Section immunostaining

Postnatal lungs were inflation-harvested as described with minor modifications (Yang et al., 2016). Briefly, mice were anaesthetized with Avertin (T48402, Sigma) and perfused through the right ventricle with phosphate-buffered saline (PBS, pH 7.4). The trachea was cannulated and the lung was inflated with 0.5% paraformaldehyde (PFA; P6148, Sigma) in PBS at 25 cm H_2_O pressure, submersion fixed in 0.5% PFA at room temperature for 4-6 hr, and washed in PBS at 4 °C overnight. Section immunostaining was performed following published protocols with minor modifications (Alanis et al., 2014; Chang et al., 2013). Fixed lung lobes were cryoprotected in 20% sucrose in PBS containing 10% optimal cutting temperature compound (OCT; 4583, Tissue-Tek) at 4°C overnight and then embedded in OCT. OCT sections at 10 um thickness were blocked in PBS with 0.3% Triton X-100 and 5% normal donkey serum (017-000-121, Jackson ImmunoResearch) and then incubated with primary antibodies diluted in PBS with 0.3% Triton X-100 in a humidified chamber at 4 °C overnight. Sections were washed with PBS in a coplin jar for 1 hr and incubated with donkey secondary antibodies (Jackson ImmunoResearch) and 4’,6-diamidino-2-phenylindole (DAPI) diluted in PBS with 0.3% Triton X-100 at room temperature for 1 hr. After another 1 hr wash with PBS, sections were mounted with Aquamount (18606, Polysciences) and imaged on a confocal microscope (A1plus, Nikon).

### Wholemount immunostaining

This was performed following published protocols with minor modifications (Yang et al., 2016). In brief, ∼3 mm wide strips from the edge of the cranial or left lobes of postnatal lungs or whole lobes of embryonic lungs were blocked with PBS with 0.3% Triton X-100 and 5% normal donkey serum (017-000-121, Jackson ImmunoResearch) and then incubated with primary antibodies diluted in PBS with 0.3% Triton X-100 overnight at 4 °C. The next day, the strips were washed with PBS+1% Triton X-100+1% Tween-20 (PBSTT) on a rocker at room temperature for one hour, and the process was repeated three times. Secondary antibodies and DAPI were added and incubated overnight at 4 °C. On the third day, the strips were washed as described before with PBSTT and fixed with PBS with 2% PFA for at least 2 hr on a rocker. For tissues expressing green or red fluorescent protein, native fluorescence was quenched after immunostaining by overnight incubation with methanol containing 6% hydrogen peroxide (H1009, Sigma) at 4 °C. Finally, the strips were mounted on slides using Aquamount (18606, Polysciences) with the flat side facing the coverslip. Embryonic lungs were also immunostained as a whole and imaged with an optical projection tomography microscope (Bioptonics, UK) as published (Alanis et al., 2014; Chang et al., 2013). Enucleated eyes were fixed in 0.5% PFA for 3-6 hr at room temperature and then retinas were dissected free of retinal pigmented epithelium, lens, and hyaloid vessels, immunostained in the same tube with matching lung strips, and mounted with the vitreous side facing the coverslip. Z-stack images of 20-40 um thick at 1 um step size were taken from the top of the tissue to obtain an en face view.

### Section in situ hybridization

Postnatal lungs were harvested and processed as described for immunostaining except 0.5% PFA was included for the sucrose/OCT overnight incubation to minimize RNA degradation. Colorimetric section in situ hybridization was carried out following published protocols (Alanis et al., 2014; Chang et al., 2013). The entire exon 3 of *Vegfa* was amplified with the following primers for probe: 5’-TGATCAAGTTCATGGATGTC-3’ and 5’-agcttataatacgactcactatagggCTGCATGGTGATGTTGCTCT-3’ (lower case indicates the T7 promoter sequence). Images were acquired on an upright Olympus BX60 microscope.

### Vasculature analysis

Wholemount samples were imaged on a confocal microscope (A1plus, Nikon) using the 40x oil objective with a field size of 318 μm x 318 μm x 20 μm and a pixel dimension of 512 x 512 x 20. At least three confocal Z-stacks per lung were analyzed with Imaris software (Bitplane) to obtain the EC number (ERG) and the percentage of proliferating (KI67) ECs. Surface rendering ICAM2 staining was used to measure vessel volume using the automatic threshold and a cut-off of 50 voxels for both control and mutant lungs. For automatic analysis of CAR4 and PLVAP staining, ERG surface rendering was filtered by the mean intensity of CAR4 or PLVAP, which was then used to mask ERG staining. To measure the EC perimeter, the viewpoint was set for the largest projection area and then all visible cell projections were measured. Cells in contact were split midway between them.

### D_2_ air space analysis and mean linear intercept analysis

The D_2_ index is an area-weighted alveolar diameter that takes into account the heterogeneous distribution of airspace sizes in the mouse lung (Jacob et al., 2009; Parameswaran et al., 2006). In this study, the D_2_ was measured and calculated on 5 um-thick frozen lung sections stained with hematoxylin and eosin (H&E). For each mouse, three images were acquired on an upright Olympus BX60 microscope with a 10x objective. Airway and main vessels that could not be avoided during imaging were filled in manually using ImageJ prior to analysis, which was performed using a 225-pixel intensity threshold and a 400-pixel area size cut-off. The same images were used to quantify the mean linear intercept with Photoshop based on a published protocol (Liu et al., 2017). Two horizontal and two vertical gridlines were drawn evenly-spaced on each picture. The distance from one alveolar wall to the next along the gridlines was measured with the Photoshop ruler tool. Airways and main vessels were excluded, as well as alveoli where one wall was not visible in the image. At least 32 intercepts per image and 3 images per mouse were measured.

### Cell dissociation and FACS

Postnatal mouse lungs were dissected in PBS, minced into pieces with forceps and digested in RPMI (ThermoFisher, 11875093) with 2 mg/mL Collagenase Type I (Worthington, CLS-1, LS004197), 2 mg/mL mL Elastase (Worthington, ESL, LS002294), and 0.5 mg/mL DNase I (Worthington, D, LS002007) for 30 min at 37 °C. The tissue was mechanically triturated after 15 min of digestion. Fetal bovine serum (FBS, Invitrogen, 10082-139) was added to a final concentration of 20% and the tissue was triturated until homogenous. The sample was transferred to the cold room and kept on ice, filtered with a 70 μm cell strainer (Falcon, 352350), and spun down at 5000 rpm for 1 min. The cells were resuspended in red blood cell lysis buffer (15 mM NH_4_Cl, 12 mM NaHCO_3_, 0.1 mM EDTA, pH 8.0) for 3 min, washed with RPMI with 10% FBS and filtered into a 5 ml glass tube with a cell strainer cap (Falcon, 352235). The cells were then incubated with CD45-PE/Cy7 (BioLegend, 103114), ICAM2-A647 (Invitrogen, A15452), CD31-BV786 (BD Biosciences, 740870) at a concentration of 1:250 for 30 minutes, spun down at 5000 rpm for 1 min, washed for 5 minutes and resuspended with RPMI with 10% FBS. The sample was refiltered and incubated with SYTOX Blue (Invitrogen, S34857), then sorted on a BD FACSAria Fusion Cell Sorter. After exclusion of dead cells, CD45 negative cells were selected and from those ICAM2 positive cells were collected.

### RT-PCR and bulk RNA-seq

RNA was extracted from FACS-purified ECs using Trizol reagents (Invitrogen, 15596018) and the RNeasy Micro kit (Qiagen, 74004). 100 ng of RNA was used for RT-PCR with the SuperScript™ IV First-Strand Synthesis System (Invitrogen, 18091050). For bulk RNA-seq, 100-200 ng total RNA was used to prepare an RNAseq library using an mRNA isolation kit (New England BioLabs, E7490) and a NEBNext Ultra RNA library prep kit (New England BioLabs, E7530S) with a final double (0.65 x −1 x bead volume) size selection step using a SPRIselect reagent kit (Beckman Coulter, B23318). The libraries were indexed (New England BioLabs, E7335S) and sequenced on an Illumina NextSeq500. Seventy-six nucleotide pair-end reads were generated for each sample and aligned to the UCSC mm10 reference genome using tophat2 and bowtie2 in R (Kim et al., 2013; Langmead and Salzberg, 2012). Transcript abundance, differential expression, and isoform quantitation were calculated using the cufflinks suite in R (Roberts et al., 2011a; Roberts et al., 2011b; Trapnell et al., 2013; Trapnell et al., 2012). Raw data have been deposit in GEO under the accession number GSE124325.

### Single-cell RNA-seq

FACS-purified lung ECs were processed through the Chromium Single Cell Gene Expression Solution Platform (10X Genomics) using the Chromium Single Cell 3’ Library and Gel Bead Kit in accordance with the manufacturer’s user guide (v2, rev D). The libraries were sequenced on an Illumina NextSeq500 using a 26X124 sequencing run format with 8 bp index (Read1). Chromium single-cell RNA-seq output was processed with Cell Ranger using “cellranger count” and “cellranger aggr”. Further analysis was carried out using Loupe Cell Browser (10x Genomics) or an R package Seurat https://satijalab.org/seurat/pbmc3k_tutorial.html. In brief, we first selected cells with at least 200 detected genes and filtered out cells with unique gene counts over 5000 or less than 200; no mitochondrial genes were found. Following filtering, we performed log normalization, identification of highly variable genes, scaling, PCA dimensionality reduction, and K-means clustering. Next, we identified differentially expressed genes per cluster and generated tSNE, dot, and violin plots, and heat maps. Raw data have been deposited in GEO under the accession number GSE124325.

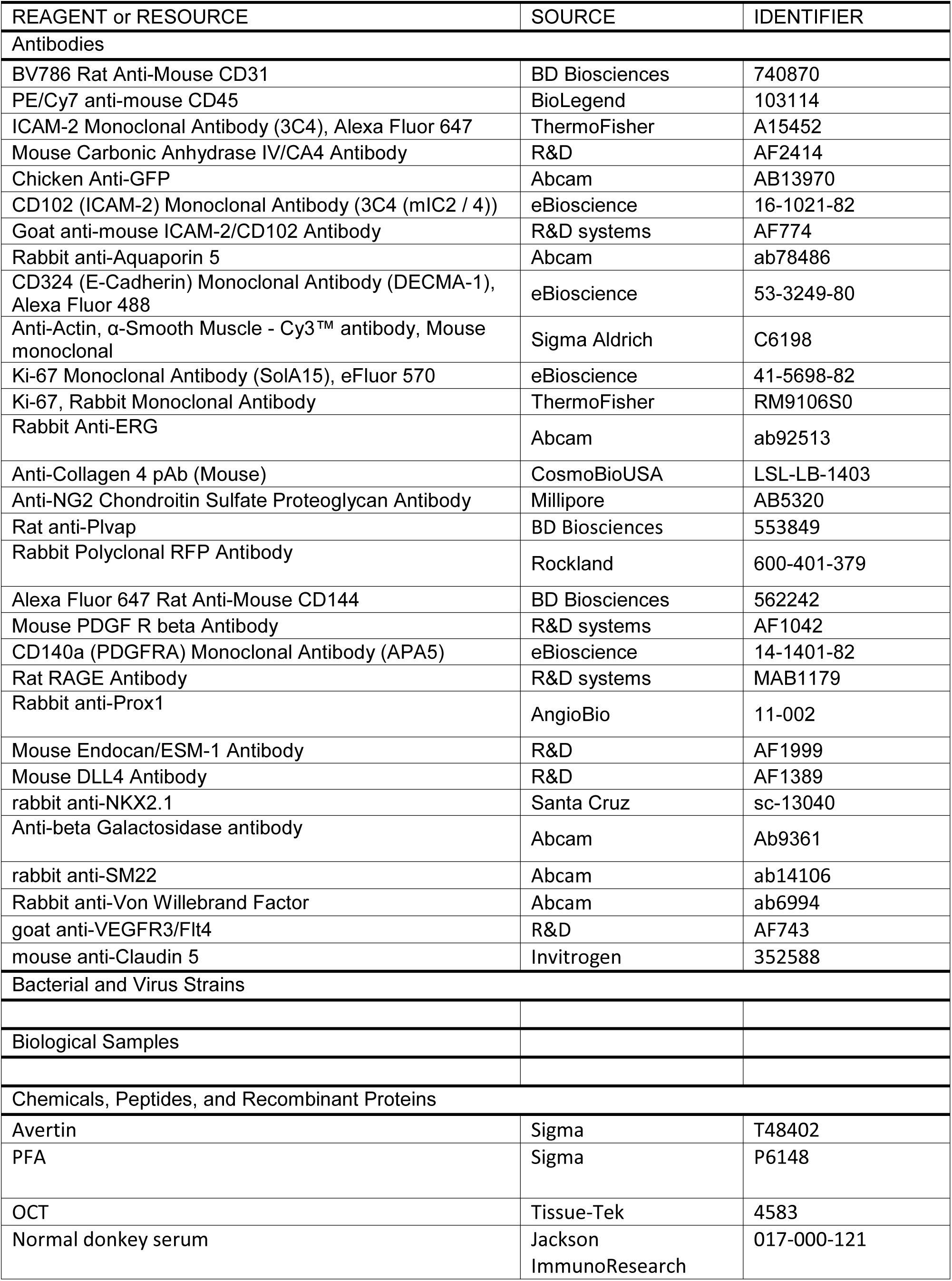
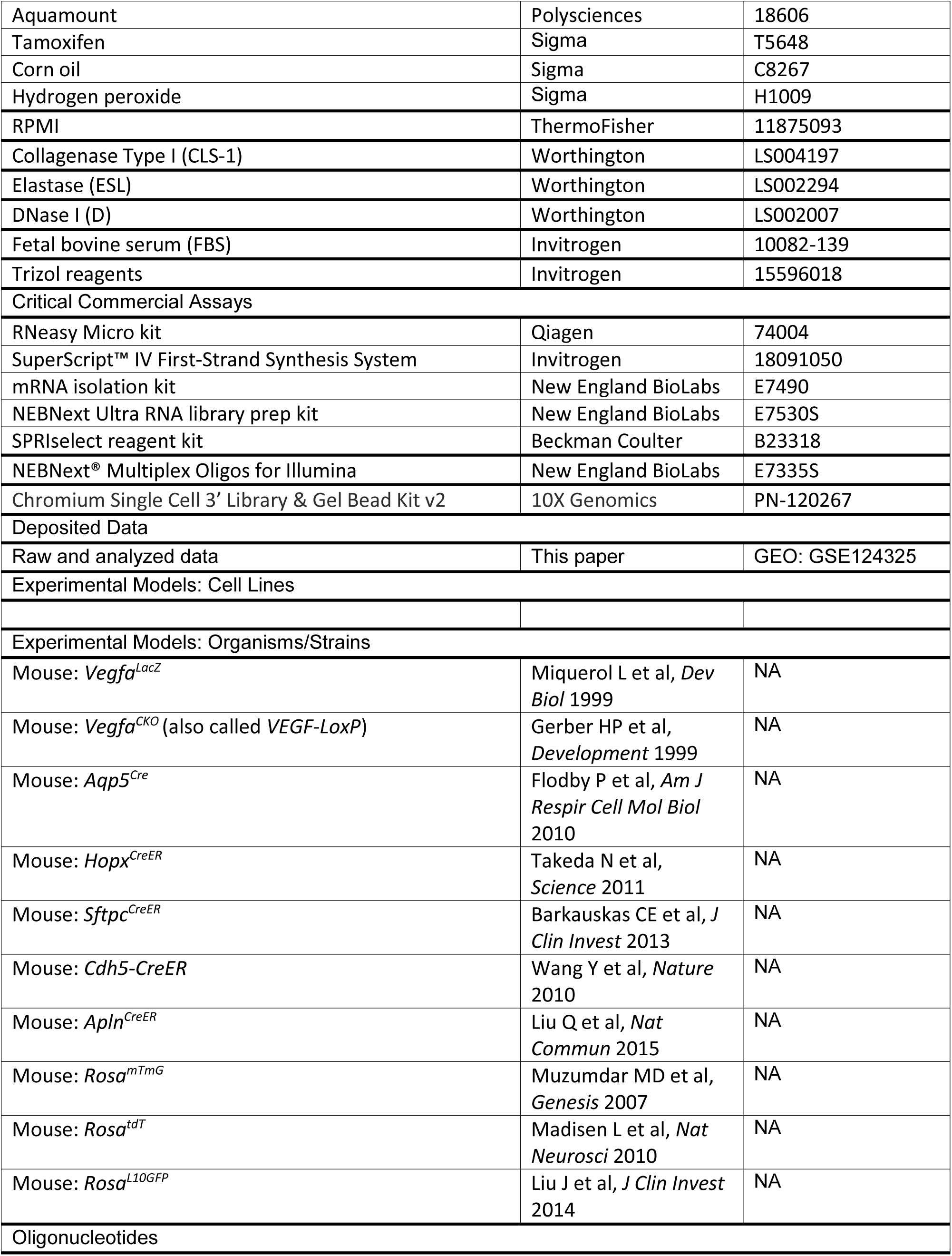
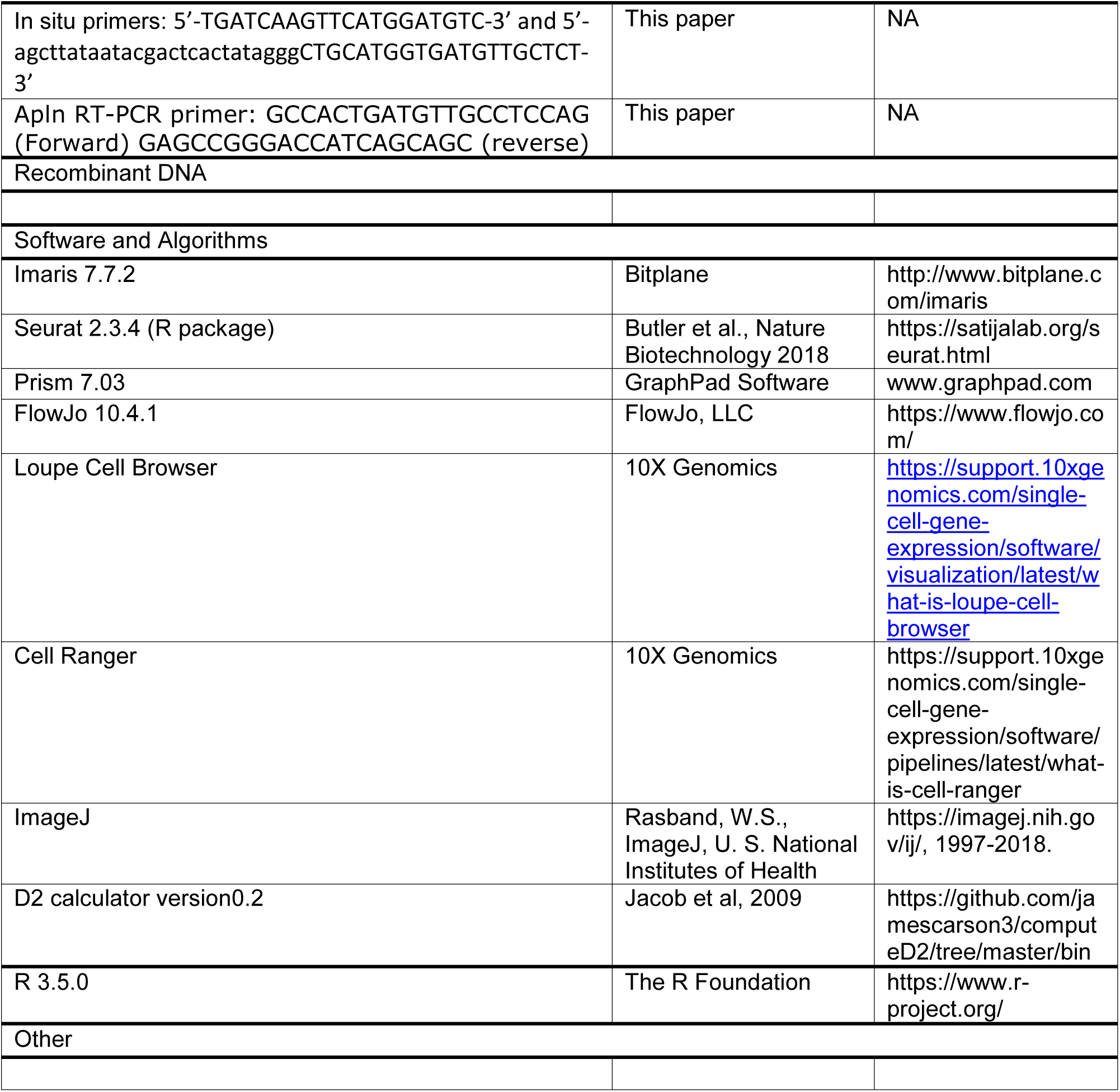

## SUPPLEMENTAL INFORMATION

**Fig. S1:**
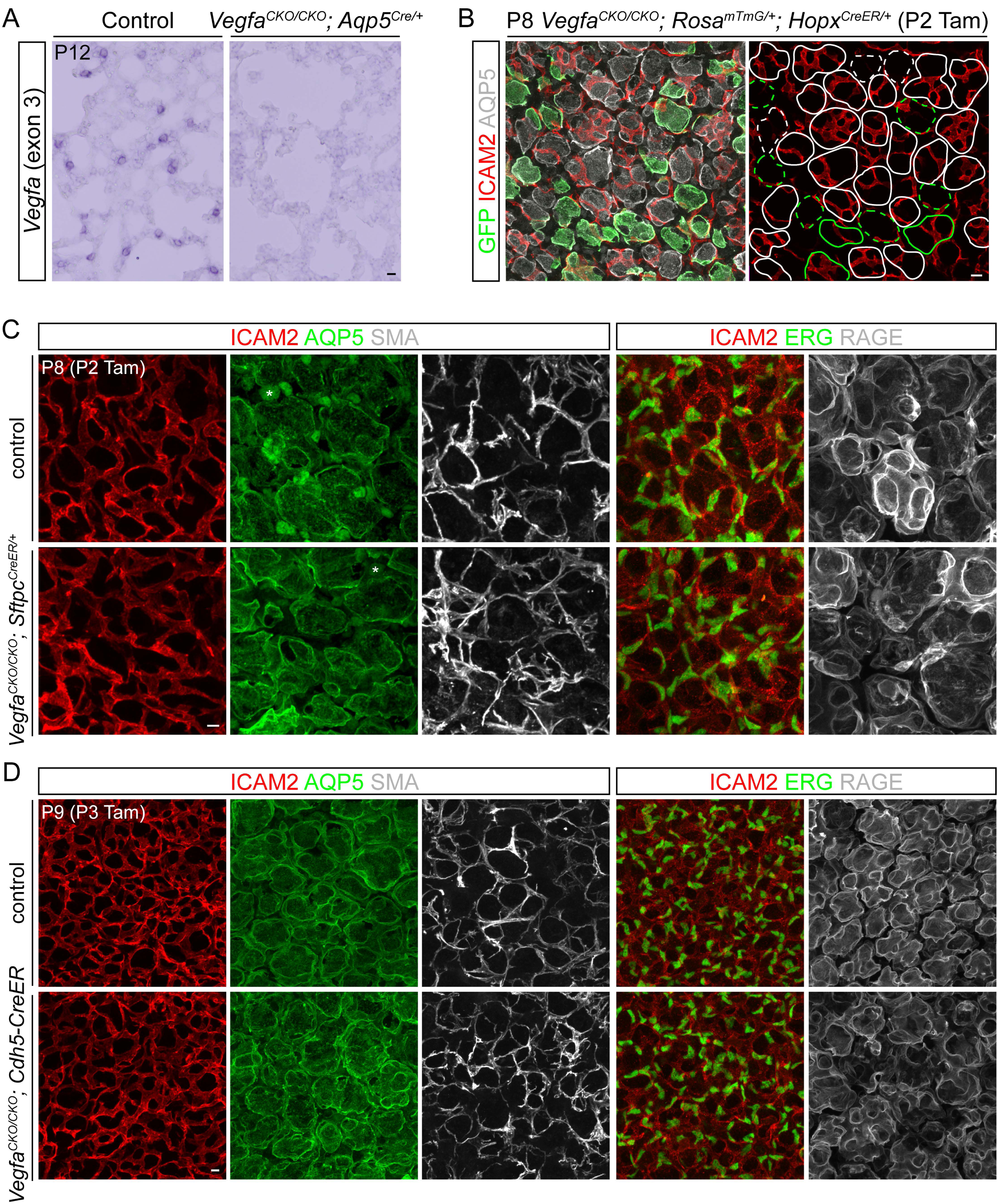
Validation of *Vegfa* deletion, quantification of mosaic deletion phenotypes, and normal angiogenesis in AT2 and endothelial *Vegfa* deletion models. Related to Figure 1, 2. (A) Section in situ hybridization of littermate lungs for the floxed exon of *Vegfa* (exon 3). Images are representative of at least 2 mice. Scale: 10 um. (B) Wholemount immunostaining to illustrate the quantification method. Alveolar islands are demarcated by examining optical sections 7 um from the top of the alveolar surface (AQP5). Islands are considered GFP^+^ (green) if >75% of the area expresses GFP. Dash, islands without vessels. Tam, 200 ug tamoxifen. Scale: 10 um. (C, D) En face view of immunostained littermate lungs, representative of at least 3 littermate pairs, showing normal angiogenesis in AT2 (C) and endothelial (D) *Vegfa* deletion models. Asterisk, non-specific nuclear staining for AQP5. Tam, 100 ug (C) and 250 ug (D) tamoxifen. Scale: 10 um.

**Fig. S2:**
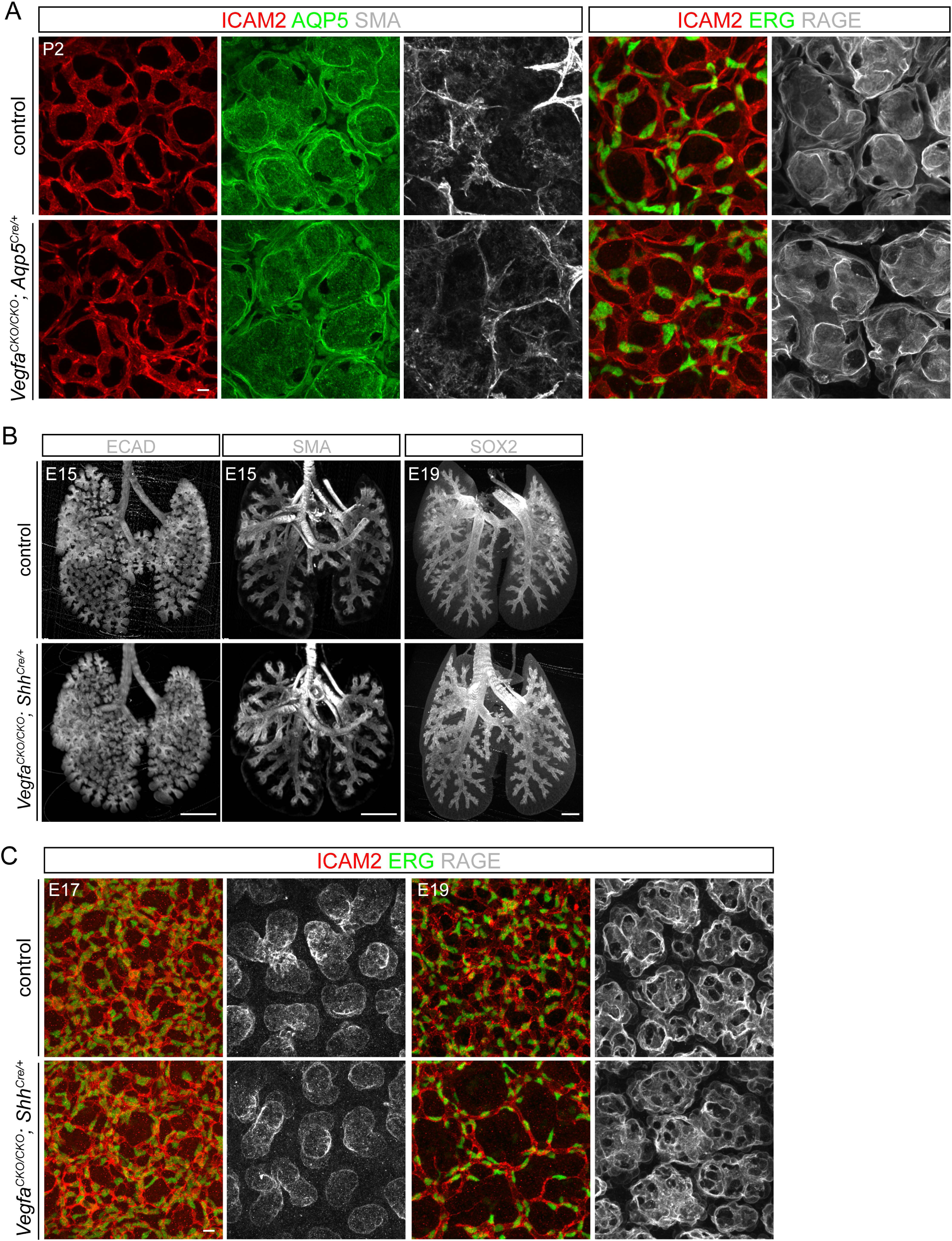
Vascular phenotypes during early development in *Aqp5^Cre^* and *Shh^Cre^* deletion models. Related to Figure 1, 3. (A) En face view of immunostained littermate lungs, representative of at least 2 littermate pairs, showing normal angiogenesis at P2. Scale: 10 um. (B) Optical projection tomography images of wholemount immunostained littermate lungs, representative of at least 2 littermate pairs, showing normal branching (ECAD), airway and vascular smooth muscle (SMA), and airway specification (SOX2). Scale: 500 um. (C) En face view of immunostained littermate lungs, representative of at least 2 littermate pairs, showing normal alveolar surface (RAGE) and sparser vasculature (ICAM2 and ERG) only after E17, concomitant with AT1 cell differentiation. Scale: 10 um.

**Fig. S3:**
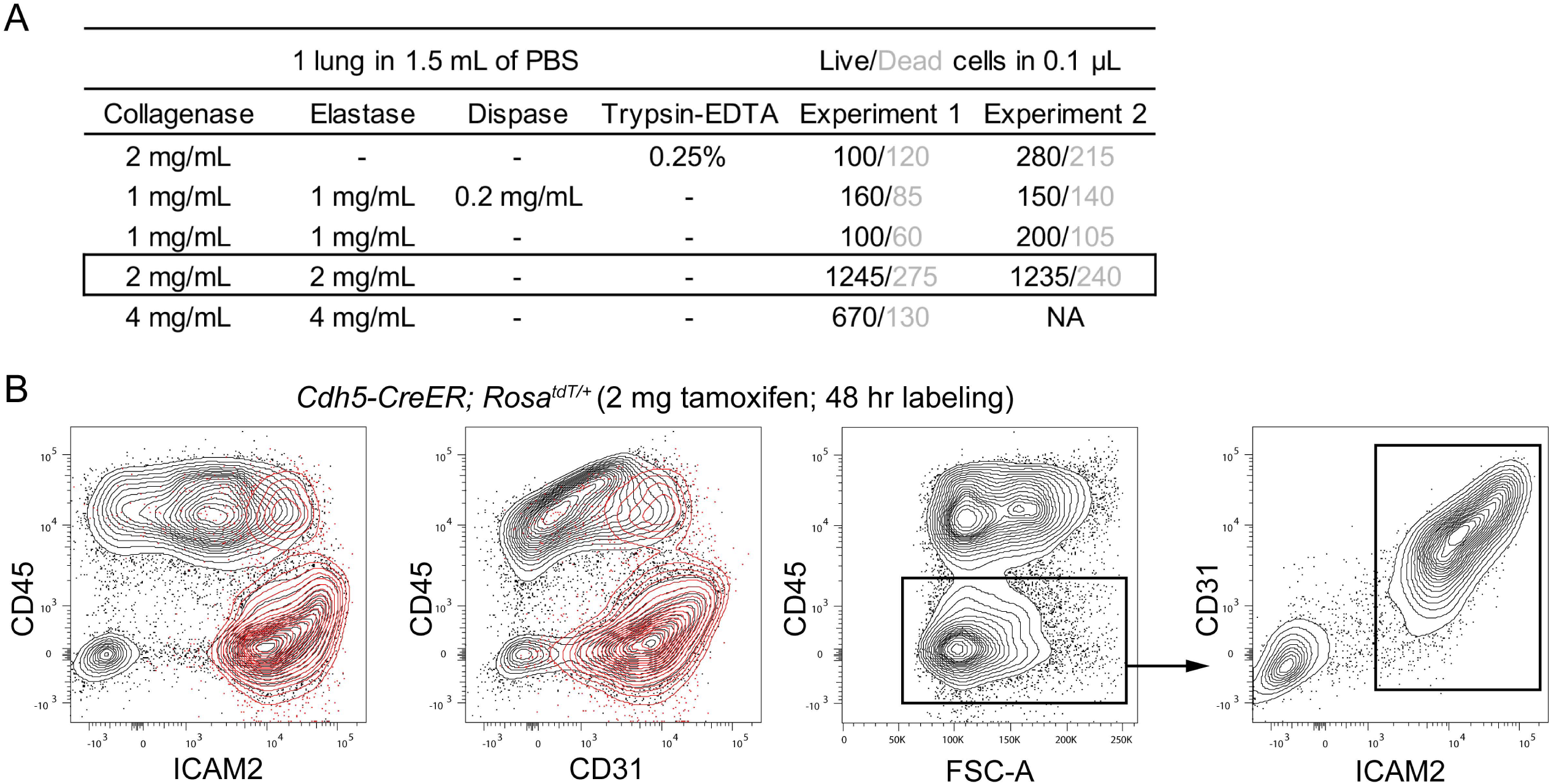
Optimization of lung EC sorting. Related to Figure 3, 4. (A) Comparison of cell dissociation protocols. For the first condition, 1.5 mL of 0.25% Trypsin-EDTA was added after collagenase digestion and the mixture was digested for an additional 15 min at 37 °C. The optimal condition (boxed) yields the highest number of viable cells in two independent experiments. NA, not available. (B) Sorting plots of an adult (>8 week-old) mouse lung with genetically labeled ECs. Left two panels: comparison of antibody staining with native fluorescence from tdT (red contour), which colocalizes mostly with CD31 and ICAM2, but also a subset of CD45 cells, perhaps due to inadvertent labeling of hematopoietic cells by *Cdh5-CreER*. Right two panels: for all sorting experiments, CD45 negative selection was first performed and followed by ICAM2 positive selection (red box), which labels the same ECs as CD31 but has better separation from the corresponding negative ECs.

**Fig. S4:**
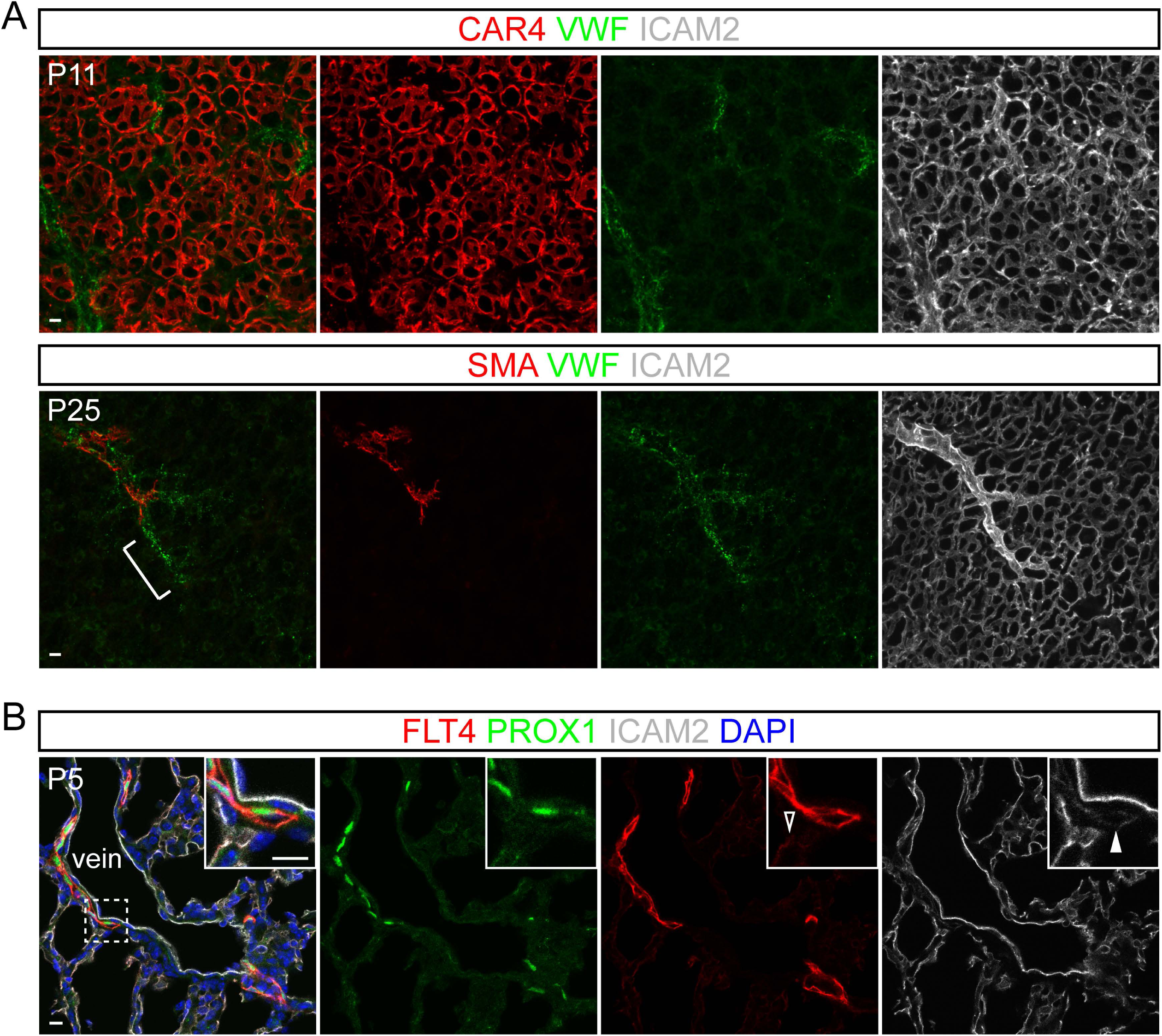
Vwf ECs and lymphatics are localized outside lung capillaries. Related to Figure 4. (A) Representative wholemount immunostaining images of at least 2 mice showing Vwf ECs in thick, non-capillary vessels (wider ICAM2 tubes) including regions just distal to SMA-wrapped vessels (bracket). Scale: 10 um. (B) Representative section immunostaining images of at least 2 mice showing lymphatic ECs (PROX1) accompanying a vein. Boxed region is magnified to show lymphatic vessels have higher FLT4 (also known as VEGFR3) but lower ICAM2 expression than blood vessels (compare open and filled arrowheads). Scale: 10 um.

**Fig. S5:**
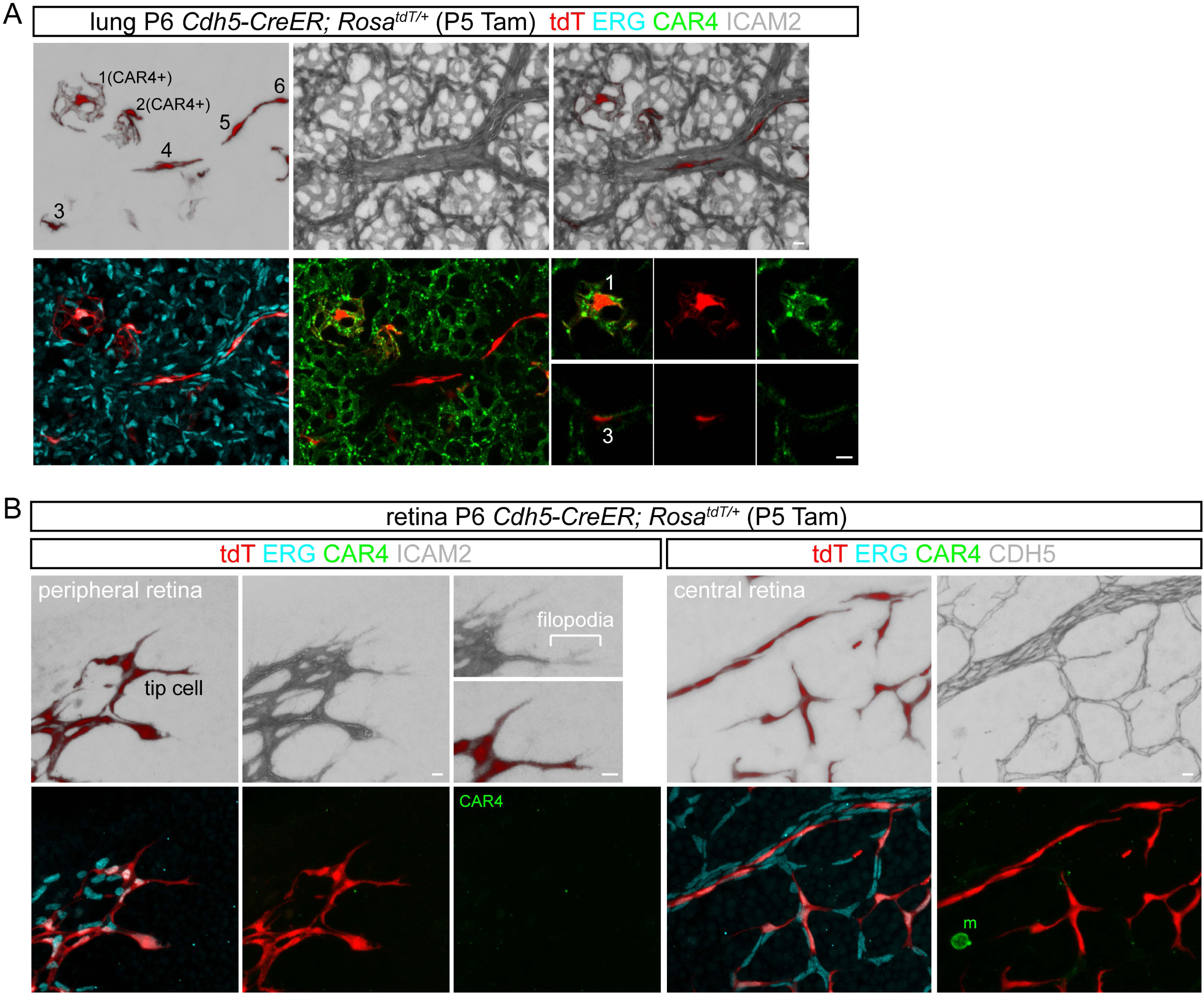
Distinct EC morphology in the lung and retina. Related to Figure 5. Wholemount immunostaining of lungs (A) and retinas (B) with sparsely-labeled ECs, representative of at least 5 mice. Tam, 0.25 ug tamoxifen. Scale: 10 um. (A) Cells #1-2 are Car4 ECs and cells #3-6 are non-Car4 ECs. ECs in non-capillary vessels (#4-6) are more elongated along the direction of blood flow. (B) Retinal macrophages (m), but not ECs, express CAR4. Tip ECs display characteristic filopodia (bracket) without a discernable lumen (weak ICAM2). Non-capillary ECs are similarly elongated, but none of the retinal ECs feature the net-like morphology of lung Car4 ECs.

**Fig. S6:**
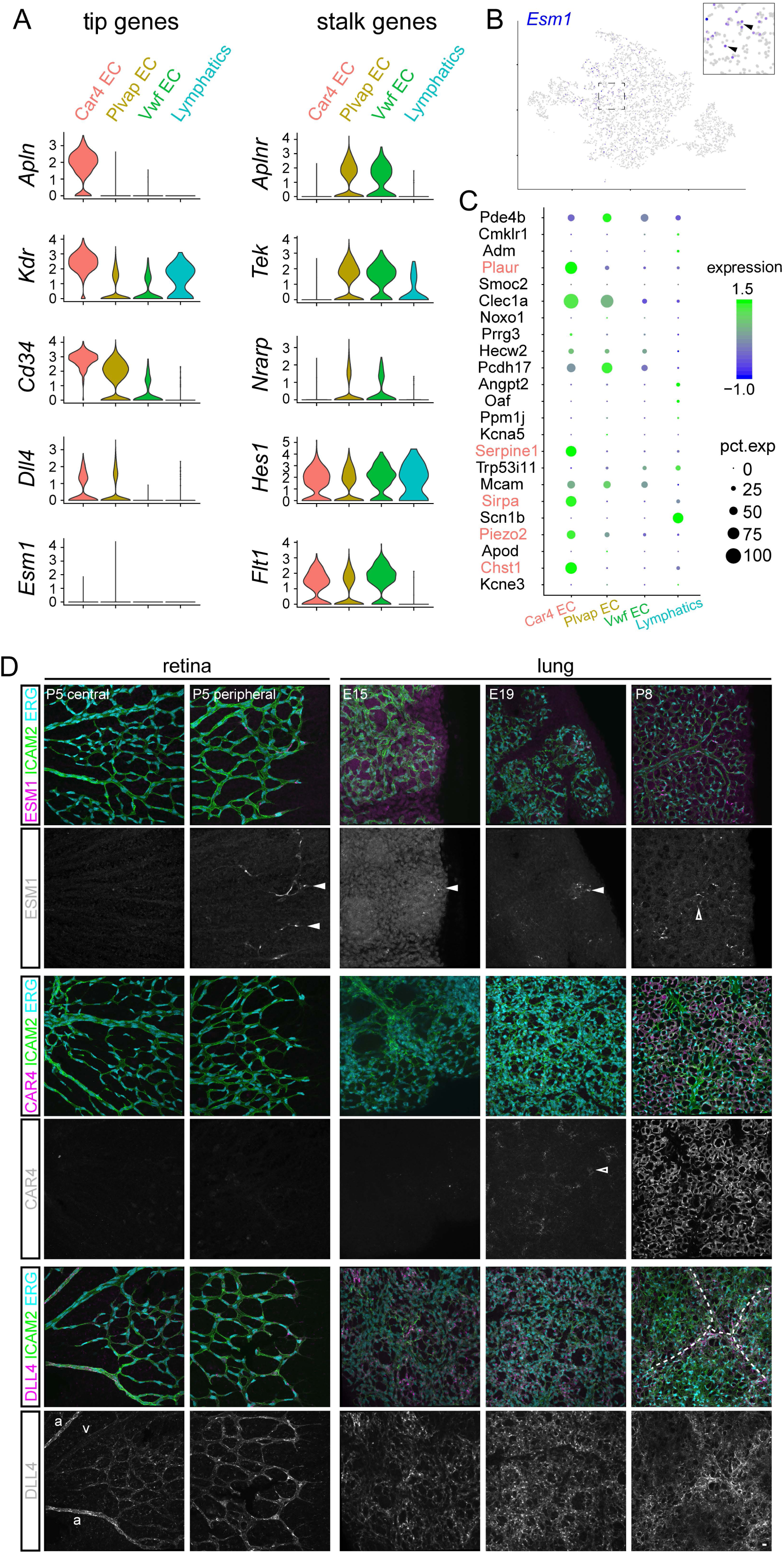
Molecular comparison of lung and retinal ECs. Related to Figure 4, 5. (A) Violin plots of lung EC scRNA-seq data from Fig. 4 showing retinal tip and stalk EC genes(Blanco and Gerhardt, 2013). *Apln* and *Kdr* are enriched in Car4 ECs; *Aplnr*, *Tek* (also known as *Tie2*), and *Nrarp* are enriched in Plvap ECs. (B) tSNE plot showing *Esm1* expression in sporadic Plvap ECs (arrowhead in insert). (C) Dot plot of 25 recently identified tip EC genes(Sabbagh et al., 2018) with those enriched in Car4 ECs highlighted in red. (D) Representative wholemount immunostaining images of at least 2 mice. ESM1 is expressed by retinal tip ECs (filled arrowhead), as well as lung ECs near lobe edge in embryos (filled arrowhead), but in transitional ECs between capillaries and non-capillaries (open arrowhead). CAR4 expression initiates at E19 (open arrowhead), concomitant with AT1 cell differentiation. DLL4 staining is stronger in retinal arteries (a) than veins (v), distinguishable by vessel diameter, and is wide-spread and cord-like (dash) in the lung. Scale: 10 um.

**Fig. S7:**
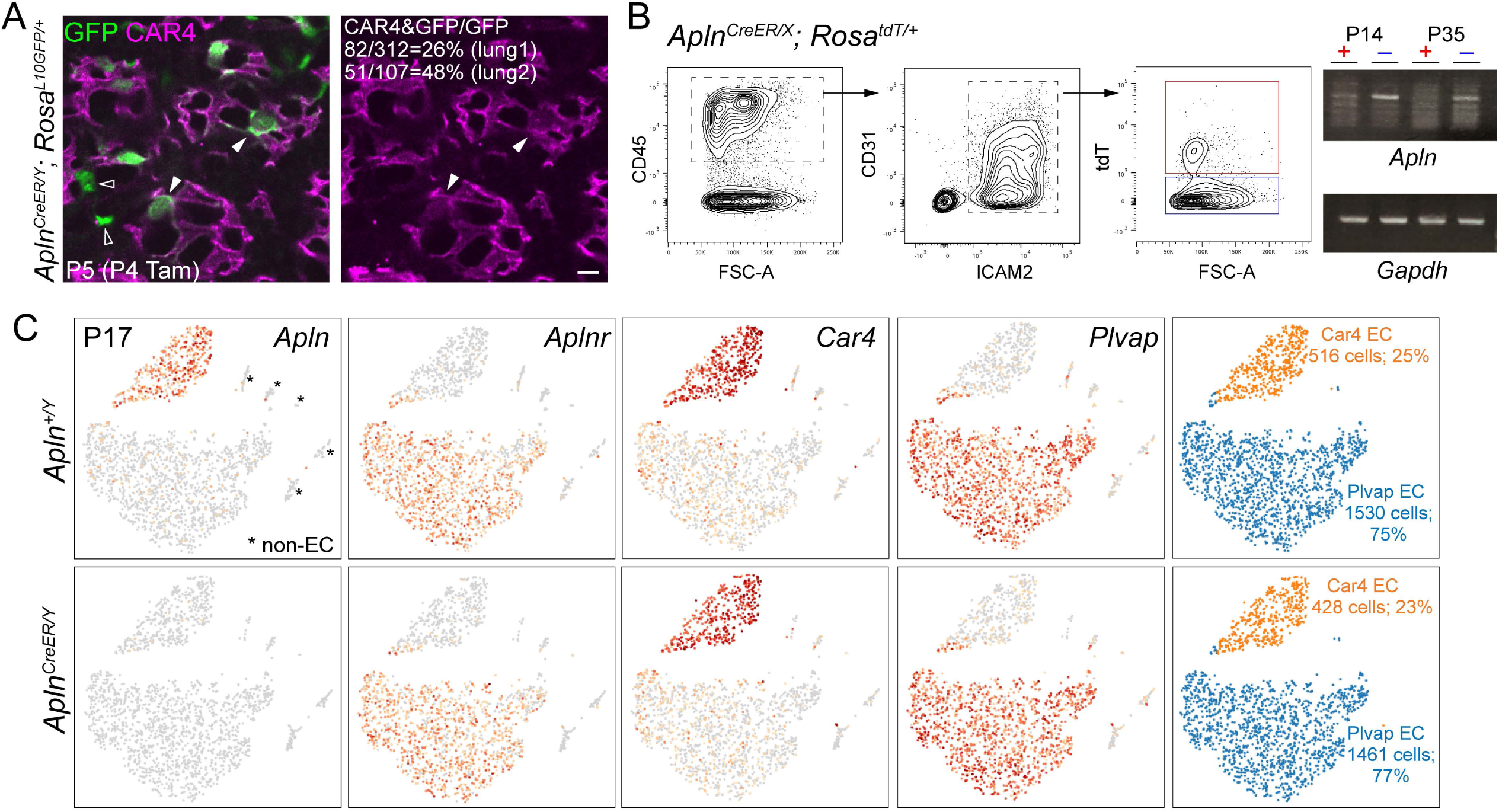
Apln mutant lungs have normal vasculature. Related to Figure 4. (A) Representative wholemount immunostaining images of at least 2 mice showing *Apln^CreER^* labels ECs but not specifically Car4 ECs. The specificity is quantified for 2 lungs and found to be variable. Filled arrowhead, CAR4 and GFP double-positive ECs; open arrowhead, GFP single positive ECs. Tam, 250 ug tamoxifen. Scale: 10 um. (B) *Apln* undergoes X-inactivation. FACS purification of P14 lung ECs (0.5 mg tamoxifen at P13) that are labeled (red box) or unlabeled (blue box) with native fluorescence from *Rosa^tdT^*, which are subjected to RT-PCR analysis. Labeled ECs, which are expected to express CreER, do not express *Apln* despite the presence of the wildtype allele, indicating X-inactivation. The same is true for P35 lung ECs (2 mg tamoxifen at P34). (C) tSNE plots of scRNA-seq results from sorted lung ECs from littermate *Apln* mutant and control males showing deletion of *Apln* but no change in the cell number and gene expression of Car4 and Plvap ECs. Asterisk, contaminating immune, epithelial, and mesenchymal cells (non-EC).

**Table S1: Bulk RNA-seq FPKM (fragments per kilobase of transcript per million mapped reads) values of sorted ECs from 3 littermate pairs of control and *Vegfa^CKO/CKO^; Shh^Cre/+^* lungs. Related to Figure 3.**

